# Prokineticin-2 Upregulates GDNF in Astrocytes and Pharmacological Modulation of PK2 Receptors offers Neuroprotection in Experimental Models of Parkinson’s Disease

**DOI:** 10.1101/2025.09.08.674934

**Authors:** Jie Luo, Griffin Clabaugh, Matthew Neal, Minhong Huang, Souvarish Sarkar, Gary Zenitsky, Huajun Jin, Vellareddy Anantharam, Canan G. Nebigil, Laurent Désaubry, Arthi Kanthasamy, Anumantha G. Kanthasamy

## Abstract

Despite a wealth of preclinical studies establishing neuroprotective and neurorestorative properties of glial cell-line-derived neurotrophic factor (GDNF) in animal models of Parkinson’s disease (PD), clinical trials utilizing direct intracranial infusion of GDNF protein, or adeno-associated virus (AAV)-mediated GDNF gene transfer has not achieved the desired efficacy, largely due to challenges in delivery methods. Given GDNF’s strong potential for neuroprotection, alternative strategies to elevate its expression by beyond invasive injection or genetic manipulation remain a promising therapeutic avenue for PD. We previously reported that prokineticin signaling provides a compensatory protective response against dopaminergic neuronal degeneration in cell and animal models of PD. Herein, we report a novel finding that PK2 regulates GDNF gene expression in astrocytes, suggesting that PK2 signaling can be harnessed for neuroprotection in PD. Treatment of cultured astrocytes with the PK2 protein, PK2 gene overexpression or prokineticin receptor 1 (PKR1) agonist IS20 significantly induced the GDNF gene expression and the protein secretion, resulting in enhanced dopaminergic cell survival in cell culture models of PD. Importantly, systemic administration of IS20 through intraperitoneal or intranasal routes elevated GDNF levels in the mouse brain, including the nigrostriatal system. Furthermore, IS20 treatment conferred significant neuroprotective effects in both 1-methyl-4-phenyl-1,2,3,6-tetrahydropyridine (MPTP)-induced and MitoPark transgenic mouse models of PD. Collectively, our translational findings suggest that pharmacological modulation PK2 signaling may unlock the full clinical benefit of GDNF, offering a novel and non-invasive therapeutic strategy for Parkinson’s disease.

## Introduction

As the second most common neurodegenerative disease, Parkinson’s disease (PD) affects around 1% of the world’s population ≥65 years old. PD is mainly characterized by progressive degeneration of dopaminergic (DAergic) neurons in the substantia nigra pars compacta (SNpc) in the ventral midbrain, resulting in dopamine depletion within the striatum. This neurodegeneration leads to hallmark motor symptoms, including bradykinesia, rigidity, and tremor^1,2^. Notably, autonomic and other non-motor deficits such as sleep disturbances, anosmia, gastrointestinal dysfunction, and mood disorders often precede the onset of motor symptoms, suggesting early involvement of extranigral brain regions in PD pathogenesis. The DA precursor, 3,4-dihydroxy-L-phenylalanine (L-DOPA), is the current standard-of-care treatment strategy for PD, which temporarily alleviates motor symptoms by increasing brain dopamine levels. However, L-DOPA has limited efficacy in addressing motor symptoms and does not halt disease progression^3–5^. Since no disease-modifying therapy has been developed, the progressive nature of the disease demands dosage titration involving increasingly higher doses of L-DOPA, resulting in an array of motor fluctuations and other severe complications, including dyskinesia^2,6^.

Glial cell-line-derived neurotrophic factor (GDNF) is well characterized neurotrophic factor involved in the development and maintenance of multiple neuronal systems including those of the kidney, enteric nervous system (ENS), and central nervous system (CNS)^7–10^. Since its discovery in 1992, GDNF has been recognized as one of the most potent neuroprotective agents, particularly within in the DAergic system^11,12^. DAergic neurons express both the GDNF family receptor GFRα1 and GDNF co-receptor RET, through which GDNF promotes neuronal survival, regeneration and functional maintenance in the SN DAergic neurons. GDNF also increases DA content and uptake, stimulates neurite extension, and increases tyrosine hydroxylase (TH) in cell culture models^12,13^. *In vivo* evidence further supports that GDNF is necessary for the homeostatic maintenance of DAergic neurons during later stages of development and throughout adulthood. Supporting this notion, conditional knockout (cKO) of GDNF in mice starting at 1-mo-old results in the selective decrease of TH immunoreactivity and a 60-70% loss of DA midbrain neurons by seven months of age, underscoring GDNF’s indispensable role in DA neuron survival under physiological conditions^10^.

Numerous preclinical studies using rodent and non-human primate models of PD have demonstrated the therapeutic potential of GDNF^14–17^. However, early clinical trials delivering recombinant (r)GDNF protein using intracranial administration failed to uniformly diffuse into deeper layers of the brain and did not slow down neurodegeneration or improve motor function^18^. While improvements in GDNF delivery have achieved moderate efficacy^19^, the difficult challenge of long-term GDNF delivery into the brain remains. Additional safety concerns including cerebellar toxicity were also evident^20^. Gene therapy approaches using adeno-associated virus (AAV)-mediated GDNF gene delivery offered the potential to overcome the delivery limitations and were deemed safe in a Phase I clinical trials. Nevertheless, Phase II trials failed to demonstrate therapeutic efficacy^21–23^. Moreover, AAV-mediated gene delivery is inherently less controllable than conventional pharmacotherapies due to irreversible integration into the host genome, raising concerns about long-term safety and regulation^24^. Failures of current GDNF trials underscore the critical need to elucidate events upstream of the GDNF signaling pathway and to disentangle the complex signaling pathways in crosstalk with GDNF. Controlling endogenous GDNF expression would constitute a powerful target in neuropharmacology and warrants increased translational research efforts^25,26^.

Setbacks of current AAV-mediated DAergic GDNF gene delivery trials force the rethinking of alternative strategies, including expressing GDNF in striatal astrocytes. Astrocytes secrete the neurotrophic factors NGF, BDNF, and GDNF^27^, and are considered primary source of GDNF in the brains of PD patients^28,29^. The role of astrocytes in maintaining a healthy environment for proper neuron function has been increasingly recognized, yet their potential as a source of nigral GNDF and as a viable therapeutic target remains underexplored. Harnessing astrocyte-mediated endogenous GDNF expression could offer a more physiologically relevant and controllable approach to neuroprotection in PD, warranting further investigation in translational research.

Initially identified from a nontoxic component of black mamba snake venom^30^, the prokineticin family proteins have been shown to interact with the GDNF signaling pathway in the peripheral nervous system (PNS) during development of the ENS^7,8,31,32^. Disruptions in either signaling pathway are implicated in Hirschsprung disease, a rare congenital disorder of the ENS^33^, highlighting the functional interdependence of GDNF and prokineticin signaling. The well-established crosstalk between these pathways in the ENS raises the intriguing possibility that the prokineticin signaling pathway also engages in similar crosstalk with GDNF in the CNS. Interestingly, both GDNF and prokineticin-2 (PK2) have been identified as potent chemoattractants for axons during neurogenesis/neuritogenesis. PK2, in particular, is essential for the complete development of olfactory bulb (OB)^34^. Importantly, our previous work demonstrated that PK2 upregulation mediates a protective, compensatory response during neurodegeneration by activating the AKT and ERK pathways and enhancing mitochondrial biogenesis^35^. AAV2/5-mediated delivery of PK2 conferred neuroprotection against Parkinsonian toxicants in the SN and striatum^35^. However, evidence is lacking for a fundamental relationship between PK2/PKR1 and GDNF/RET signaling in the CNS and the nigrostriatal system. Intriguingly, PKR1 is mainly expressed in astrocytes^36^, raising the possibility that GDNF may be co-upregulated with the prokineticin pathway in astrocytes. The preferential expression of PKR1 over PKR2 in astrocytes further supports the feasibility of selectively targeting astrocytes using PKR1-specific agonists. Based on this therapeutic rationale, using IS20, a lipophilic small molecule receptor agonist specific to PKR1^37^, we hypothesized that positive modulation of the expression and secretion of GDNF in astrocytes of the nigrostriatal system could be achieved via pharmacological activation of PK2/PKR1 signaling. This astrocyte-derived trophic support may, in turn, promote the DAergic neuronal survival and exert both neuroprotective and neurorestorative effects in experimental models of PD.

In this study, we report that 1) PK2/PKR1 signaling represents a novel, druggable pathway for modulating GDNF expression in the CNS, particularly in astrocytes, 2) the upregulation of GDNF through PK2/PKR1 signaling can be pharmacologically activated by intranasal (IN) or intraperitoneal delivery of the PKR1 receptor agonist IS20, and 3) IS20 could rescue DA neurons and neuronal projections from neurodegeneration in genetic and chemically-induced mouse models of PD. Unlike the uncontrolled, long-term continuous expression driven by intracerebral injections of GDNF viral vectors, which leads to unwanted compensatory responses^38–40^, systemic administration small molecule IS20 showed no such adverse effects, nor any overt toxicity in mice. This distinction highlights advantage of pharmacologically modulating endogenous GDNF expression more controlled and physiologically relevant manner. Thus, by pharmacologically modulating the magnitude and duration of endogenous GDNF expression using small-molecule IS20 treatment, GDNF’s established clinical benefits could be fully harnessed.

## Results

### Prokineticin receptor 1 is activated by rPK2 protein and receptor agonist IS20

Previous work in our lab has demonstrated that PK2 signaling functions as a potent neuroprotective response during early stages of DAergic neurodegeneration in models of PD. Given the established crosstalk between prokineticin signaling and GDNF during development, and the well-characterized protective effect of GDNF in preclinical models of PD, we hypothesized that PK2’s neuroprotective functionalities are afforded through activation of GDNF signaling. PKR1 is expressed on the membrane of DAergic neurons and Type 1 astrocytes in the nigrostriatal system. We confirmed PKR1 and GDNF expression in the substantia nigra by immunostaining human postmortem nigral sections probing for PKR1 and GDNF (Supplementary Fig. 1a). Interestingly, we found that PKR1 is highly expressed in the SN and colocalized with GDNF, indicating the possibility of GDNF expression being downstream of PKR1 signaling. We also found a marked decrease in GDNF and mRNA (Supplementary Fig. 1b) in PD patient nigral lysates compared to control samples in line with some previous reports^29,41^ while conflicting with others^42^. This decrease in GDNF highlights the lack of trophic support in the nigrostriatal system in PD patients and the therapeutic potential of restoring GDNF levels in the SN.

To examine whether the PK2 agonist IS20 can positively modulate GDNF expression, our initial set of experiments was conducted to confirm that IS20 activates PKR1 as has been previously reported^37,43^. Once activated, PKR1 initiates downstream signaling that culminates in the rise of Ca^2+^ in the cytosol^37,44^. *In silico* modeling with AlphaFold3 of PK2 binding to PKR1 confirms that the AVITGA sequence motif resides in the binding pocket of PKR1 (Supplementary Fig. 2a). This is in alignment with previous reports of AVITGA being required for receptor binding. In silico analysis also revealed that IS20 can bind to PKR1 with high affinity (Supplementary Fig. 2b).

To examine PKR1 activation *in vitro*, we used a CHO cell line stably expressing PKR1. The PKR1-overexpressing cells were treated with 100 nM rPK2 or 10 µM IS20, and PKR1 activation was monitored by measuring intracellular Ca^2+^ mobilization in real time over 180 s using a Fluo-4 NW calcium assay kit. Upon treatment with rPK2, intracellular Ca^2+^ concentration rose rapidly, peaking at 20 s post-treatment (Supplementary Fig. 2c), indicative of a Gαq1 mechanism for PKR1 activation. Treatment with 10 µM of IS20 caused Ca^2+^ concentration to rise more slowly, peaking at 180 s post-treatment (Supplementary Fig. 2d), suggesting that IS20 induces a similarly large but relatively delayed activation of PKR1 compared to rPK2.

### Recombinant PK2 and PK2 agonist IS20 upregulate GDNF expression in human astrocytes

To examine the effects of PK2 signaling activation on GDNF expression in astrocytes, we first incubated U373 human astrocyte cells with rPK2 protein dissolved in reduced serum (2% FBS). Treatment with 25 nM rPK2 for 8 h induced GDNF mRNA expression levels to nearly 2.5-fold above control (Fig. 1a), and treatment for 10 h induced GDNF protein levels 1.7-fold above control (Fig. 1b). The rPK2-induced upregulation of GDNF mRNA was completely attenuated by co-treatment with the prokineticin receptor antagonist PC-7 (Fig. 1a). To validate these findings in human cells, primary human astrocytes were incubated with 25 nM rPK2 for 4, 8, and 12 h. We found that rPK2 incubation elicited a time-dependent increase in GDNF mRNA over 8 h that diminished by 12 h (Fig. 1c). Immunocytochemistry (ICC) analysis confirmed the increase in GDNF protein levels compared to control (Fig. 1d).

**Figure 1.**
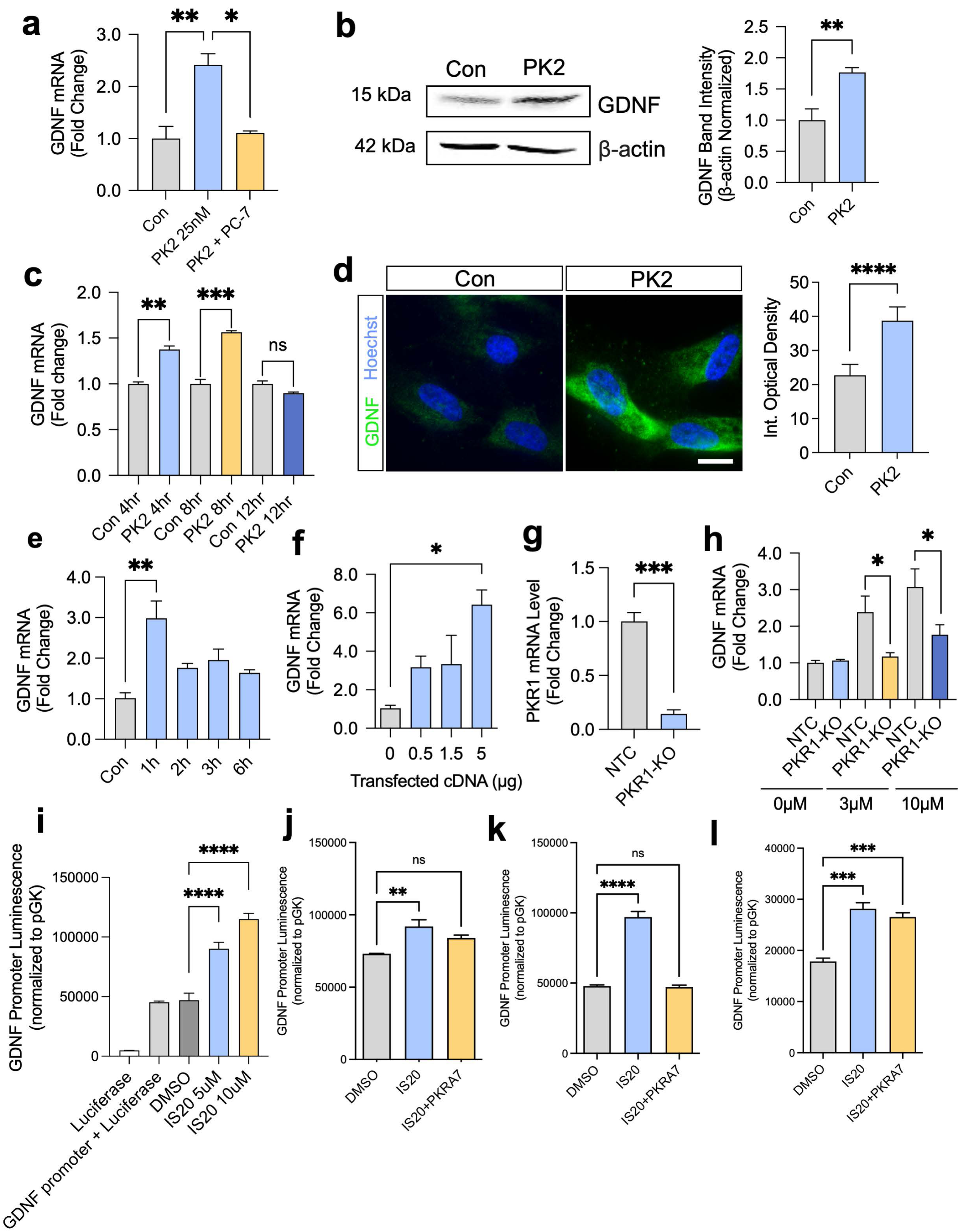
PK2 and PK2 agonist IS20 induce GDNF expression in human astrocytes. (a) qPCR of GDNF gene expression in U373 human astroglioma cell line treated with PK2 (25 nM) or co-treated with PK2 (25 nM) and 1 µM PC-7, a PKR1/PKR2 inhibitor, for 8 h (n = 4). (b) Western blot analysis probing for GDNF in U373 cells treated with PK2 (25 nM) for 10 h (n = 4). (c) qPCR analysis of GDNF mRNA from primary human astrocytes treated with PK2 for 4, 8, and 12 h (n = 2). (d) Immunocytochemical analysis probing for GDNF in primary human astrocytes treated with 25 nM PK2 for 10 h (n = 8; scale bar = 10 µm). (e) qPCR analysis of GDNF gene expression of U373 cells treated with IS20 (10 µM) for 1-6 h (n = 3). (f) GDNF mRNA levels measured by qPCR of U373 astrocytes transiently transfected with a PKR1 overexpression vector with increasing amounts of cDNA (0.5–5 µg) and treated with 10 µM IS20 for 4 h (n = 3–4). (g) qPCR analysis of PKR1 mRNA confirming CRISPR-Cas9 PKR1-KD. (h) qPCR analysis of GDNF mRNA in CRISPR-Cas9 PKR1-KD cells treated with IS20 (3–10 µM) for 8 h (n = 6). (i) GDNF promoter luminescence assay of U373 astrocytes transfected with 3.6 kb of GDNF promoter fragment and 24 h after transfection, treated with IS20 (5 µM or 10 µM) for 8 h. (j) GDNF promoter luminescence following 4 h, (k) 8h, and (l) 12h of 10 µM IS20 in U373 astrocytes transfected with GDNF promoter fragment. *p≤0.05, **p<0.01, ***p<0.001, ****<0.0001.

Next, we sought to determine if IS20 pharmacological activation of PKR1 could modulate GDNF in U373 astrocytes. A time-course treatment of U373 cells with 10 µM IS20 revealed a 3-fold increase in GDNF mRNA after 1 h of IS20 incubation; this upregulation declined over the subsequent 5 h (Fig. 1e). To assess any receptor-specific effects of IS20 on GDNF production, we overexpressed PKR1 by transiently transfecting increasing amounts of PKR1 expression vector in U373 astrocytes. Ectopic expression of PKR1 (0.5–5 µg cDNA) dose-dependently stimulated the upregulation of GDNF mRNA in cells treated with 10 µM IS20 for 4 h (Fig. 1f). Importantly, when a CRISPR-Cas9 guide RNA against PKR1 was used to knock down PKR1 in U373 astrocytes (Fig. 1g), IS20 was no longer able to modulate GDNF expression (Fig. 1h), further demonstrating that IS20-induced GDNF expression is highly dependent on PKR1 receptor binding and activation.

To validate our findings that IS20 could enhance GDNF expression and increase GDNF promoter activity, we utilized a GDNF promoter luciferase assay. U373 astrocytes were transfected with a 3.6 kb GDNF promoter fragment that encompasses GDNF promoter 1, TATA-box, the main transcriptional start site and exon 1 of the murine GDNF gene cloned into a pGL3 promoter vector that drives the expression of a luciferase reporter gene. 24 h after transfection, U373 cells were treated with 5 µM and 10 µM of IS20 for 8 h. We observed a dose-dependent increase in luciferase signal intensity with both doses significantly enhancing GDNF promoter activity (Fig. 1i). Next, we treated the transfected U373 cells with IS20 (10 µM) for 4, 8, and 12h. Notably, we observed a significant increase in relative luciferase signal intensity at all timepoints. Co-treatment of PKR1 antagonist, PKRA7, with IS20 attenuated the increase in the 4h and 8h groups while the reduction in GDNF promoter activity was not seen in the 12h group. This could be due to PKRA7 degradation after 12 h leaving it ineffective at blocking PKR1 activation. Together, these results suggest that IS20 can enhance GDNF promoter activity and gene expression in U373 astrocytes.

### GDNF is upregulated in primary mouse astrocytes and organotypic midbrain slice cultures treated with rPK2 and IS20

Next, we used cultured primary mouse astrocytes from prenatal mouse pups to validate the findings obtained from the human astrocyte cells. Treating primary mouse astrocytes with rPK2 (10–25 nM) for 8 h revealed a 3.4-fold upregulation of GDNF mRNA in the 25-nM group, which was abolished with PC-7 antagonism (Fig. 2a). Western blot and ELISA analysis confirmed a corresponding 1.5-fold increase in PK2 protein levels (Fig. 2b–c). Similarly, using primary mouse astrocytes stably expressing PK2 tagged with the green fluorescent protein (GFP) reporter (Fig. 2d), we found Lenti-PK2-GFP cells exhibited a 1.7-fold increase in GDNF protein levels compared to Lenti-GFP-infected control cells (Fig. 2e).

**Figure. 2.**
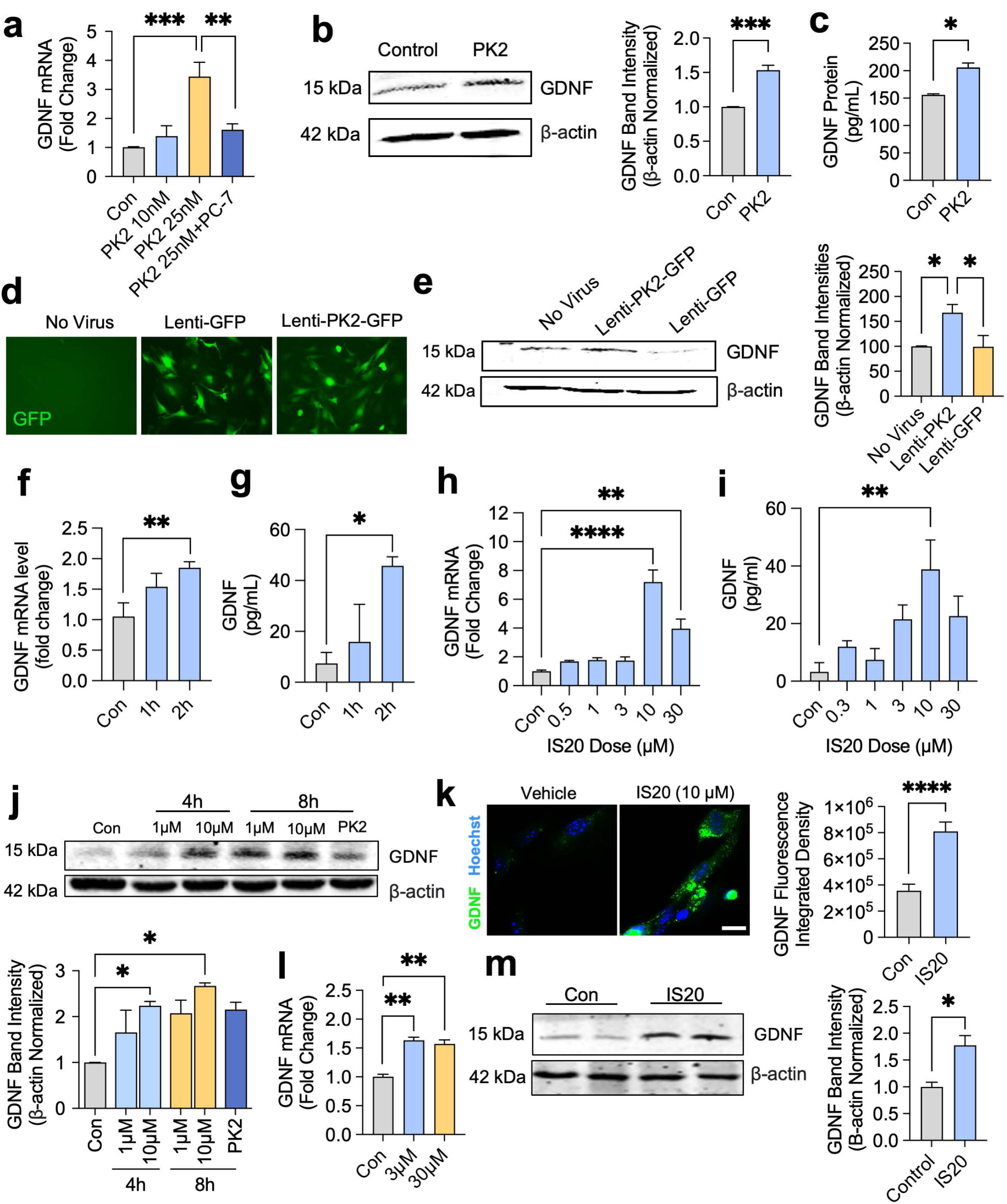
PK2 and IS20 increase GDNF in primary cultures of mouse astrocytes and organotypic slices. (a) qPCR analysis of GDNF mRNA in primary mouse astrocytes treated with PK2 (10–25 nM) or co-treated with PK2 and PC7 (1 µM) for 8 h (n = 6). (b) Western blot analysis probing for GDNF in primary mouse astrocytes treated with PK2 (25 nM) for 10 h (n = 3–5). (c) ELISA assay of GDNF protein from mouse primary astrocyte lysates treated with 25 nM PK2 for 10 h (n = 2). (d) Fluorescence microscopy showing stable expression of PK2-eGFP after lentiviral-mediated delivery of PK2 cDNA into primary mouse astrocytes. (e) Western blot analysis of GDNF protein level in primary mouse astrocytes transduced by lentiviral PK2-eGFP (n = 4). (f) qPCR assay of GDNF mRNA in primary mouse astrocytes treated with 10 µM IS20 for 1–2 h (n = 3). (g) GDNF ELISA of the supernatant of primary mouse astrocytes treated with 10 µM IS20 for 1–2 h (n = 3). (h) qPCR analysis of GDNF mRNA of primary mouse astrocytes treated with 0.5–30 µM IS20 for 3 h (n = 3). (i) GDNF ELISA of the supernatant of primary mouse astrocytes treated with 0.3–30 µM IS20 for 2 h (n = 3). (j) Western blot analysis probing for GDNF (left) in primary astrocyte lysates treated with 1 µM or 10 µM IS20 or 25 nM PK2 for 4 h or 8 h and densitometric analysis (right) of GDNF band normalized to β-actin (n = 2). (k) GDNF immunocytochemistry of primary human astrocytes treated with IS20 (10 µM) for 8 h (n = 4, scale bar = 10 µm). (l) qPCR analysis of GDNF gene expression in cultured midbrain organotypic slices treated with 0, 3, or 30 µM IS20 for 4 h (n = 3). (n) Western blot analysis of GDNF protein in cultured organotypic slices treated with 0 or 30 µM of IS20 for 4 h (n = 3). *p≤0.05, **p<0.01, ***p<0.001, ****p<0.0001.

To evaluate the toxicity of IS20 on primary mouse astrocyte survival, we utilized the MTS assay. We found the LD50 toxicity at 24 h was between 348 µM and 592 µM IS20 (Supplementary Fig. 3a). To assess the GDNF response to IS20, we treated primary mouse astrocytes with 10 µM IS20 for 1–2 h. We found that IS20 induced rapid upregulation of GDNF mRNA, as measured by qPCR (Fig. 2f), and corresponded to a rise in GDNF protein in the media as measured by ELISA (Fig. 2g). A dose–response study was utilized to determine the optimal dose of IS20 in primary mouse astrocyte cells to elicit a GDNF response. Treating astrocyte cells with 0.5-to 30-µM IS20 for 2 h revealed that maximal GDNF expression occurred at 10 μM IS20 (Fig. 2h–i). Interestingly, mRNA of the predominant GDNF receptor in the DAergic system, GFRα1, was co-upregulated with GDNF in the 10-μM IS20 treatment group (Supplementary Fig. 3b), indicating that the increased gene expression and protein secretion of GDNF also lead to upregulated ligand–receptor interaction, possibly through a positive feedback loop, to promote GDNF signaling. GDNF protein levels were also increased in whole cell lysates treated with 1 and 10 µM of IS20 for 4-8 h (Fig. 2j). Moreover, ICC analysis confirmed the increase in GDNF following 8 h of 10 µM IS20, further demonstrating that GDNF is both upregulated and secreted by primary astrocyte cells following IS20 treatment.

To evaluate the specificity of IS20 in stimulating GDNF expression, we used IS21, an analog of IS20 incapable of activating PKR1. Treating primary mouse astrocytes with IS21 for 3 h did not confer GDNF expression, while IS20 induced a 3-fold increase in GDNF mRNA (Supplementary Fig. 2b). Moreover, using Qiagen Neurotrophin and Receptor PCR array analysis, we found increased expression of 71 out of 84 genes in primary mouse glial cultures treated with IS20, including GDNF, CTNF, and BDNF (Supplementary Fig. 2c), further suggesting IS20 can promote neurotrophic support in mouse astrocytes.

After establishing IS20’s effects on GDNF expression using astrocyte cultures, we validated those findings using an *ex vivo* approach employing organotypic midbrain slice cultures prepared from neonatal mouse pups. In line with our *in vitro* findings, treating organotypic midbrain slices with IS20 for 4 h increased GDNF mRNA expression (Fig. 2l) and protein levels (Fig. 2m).

Collectively, these results demonstrate that GDNF is highly upregulated and secreted in both primary mouse astrocyte and midbrain slice cultures following PK2 and IS20 activation of PKR1 signaling.

### IS20 protects against MPP^+^-induced cell death and preserves mitochondrial energetics in DAergic neuronal cells

Having discovered that IS20 is a potent inducer of GDNF in astrocytes, we further evaluated whether the IS20-induced upregulation and secretion of astrocytic GDNF could functionally protect DAergic neuronal cells under insult from the classic Parkinsonian toxicant MPP^+^. U373 astrocytes were incubated with or without IS20 for 2 h, followed by a change to fresh media and an additional incubation for 6 h. Both types of astrocyte-conditioned media (ACM) were collected and added to N27 DAergic neuronal cells that were then co-treated with or without MPP^+^ (100– 300 µM) for 24 h (Fig. 3a). Adding IS20-treated ACM significantly increased cell viability in MPP^+^-treated N27 cells compared to MPP^+^-treated cells that had received only vehicle-treated ACM (Fig. 3b). Next, we investigated whether this protection was mediated through preservation of mitochondrial respiration. Seahorse (Agilent Technologies, Santa Clara, CA) analysis revealed that, in addition to slightly increasing the basal respiration of untreated N27 cells (Fig. 3c), IS20-treated ACM remarkably attenuated the MPP^+^-induced reduction in oxygen consumption rate, basal respiration and reserve capacity (Fig. 3d), suggesting that a post-mitochondrial mechanism at least partly contributes to the protection conferred by IS20-treated ACM. Lastly, we utilized mouse primary mesencephalic culture to validate the neuroprotective effects of IS20-treated ACM. We included the ACM collected from U373 astrocytes treated with IS21, an analog of IS20 that does not activate PKR1, to further determine the specificity of IS20. IS20-treated, but not IS21-treated, ACM promoted cell survival in MPP^+^-exposed mouse primary mesencephalic culture as determined by MTS assay (Fig. 3e). Altogether, these studies suggest that U373s treated with IS20 secrete factors that protect DAergic neuronal cells against MPP^+^-induced neurotoxicity.

**Figure 3.**
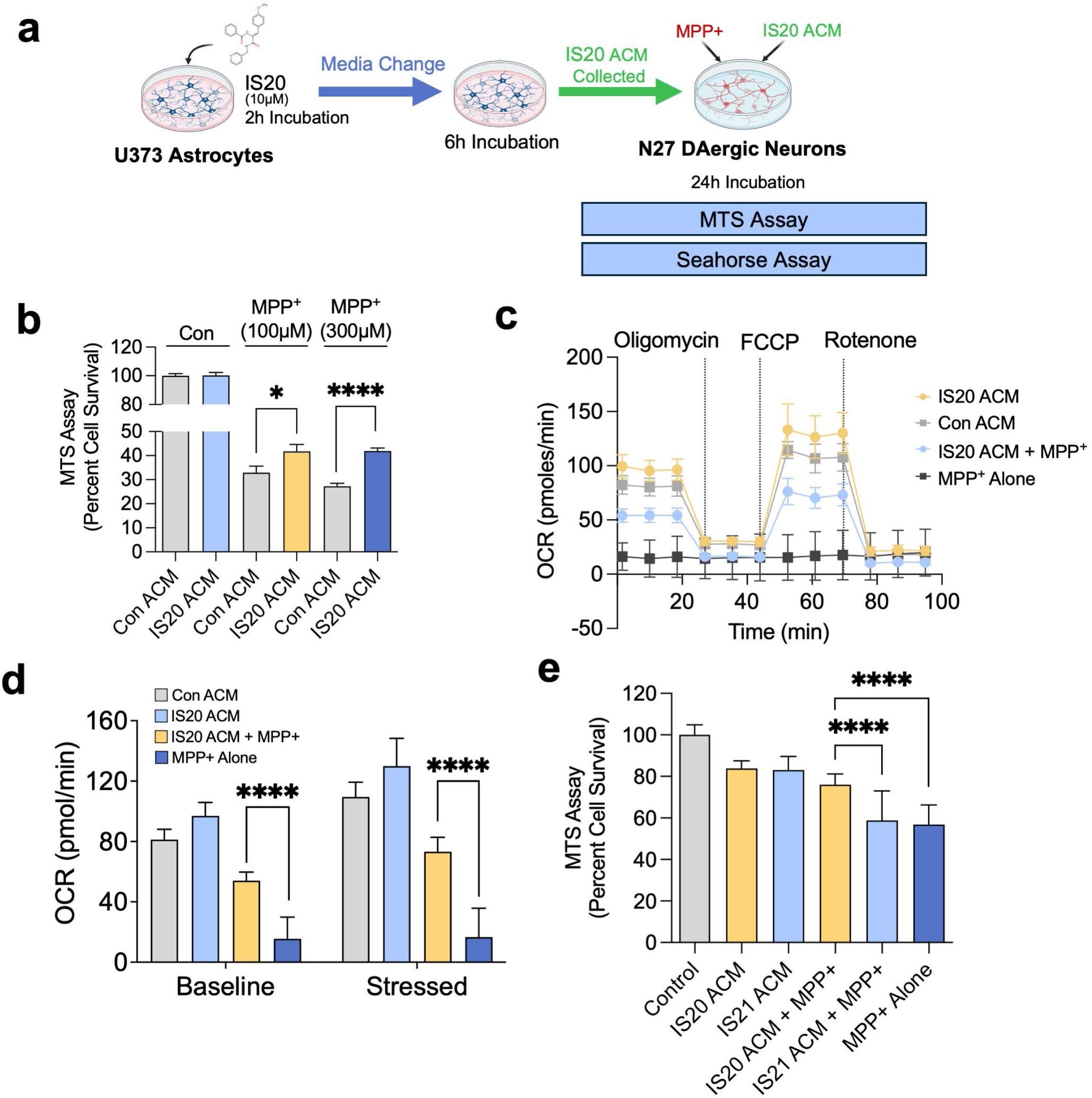
IS20 protects against MPP+-induced cell death and preserves mitochondrial energetics in DAergic neuronal cells. (a) Schematic of the IS20 treatment paradigm whereby U373 astrocytes were treated with 10 µM IS20 or vehicle control for 2 h, the media was changed and the U373 astrocyte cells were incubated for 6 h. Next, the astrocyte media was collected and added to N27 DAergic neuronal cells that were co-treated with or without MPP+ (100 µM or 300 µM). (b) MTS assay of N27 DAergic neurons after co-treatment with MPP+ and astrocyte-conditioned media (ACM) that had been incubated with or without IS20 for 24h (n = 6-7). (c–d) Seahorse Mito Stress Test of N27 cells treated with control ACM (grey line), IS20 ACM (yellow line), IS20 ACM added with MPP+ (blue line), and MPP+ alone (black line) (n = 12). (e) MTS assay of primary striatal culture isolated from the mouse and treated with the same control ACM, control ACM with MPP+, IS20 ACM + MPP+, IS21 ACM + MPP+, and the controls. *p≤0.05, **p<0.01, ***p<0.001, ****p<0.0001.

### IS20 administration induces GDNF expression and release in C57BL/6 mice

Next, we determined the bioavailability of IS20 in the brain after intraperitoneal (i.p.) injection or IN administration and assessed its biological effects in the brain. First, wild-type C57BL/6 mice were i.p.-injected with DMSO or IS20 at 10 mg/kg body weight. Mice were sacrificed after 8 h to examine GDNF protein levels in the brain and serum by Western blot and ELISA, respectively. IS20 significantly increased GDNF protein levels in both nigral lysates (Fig. 4a) and serum (Fig. 4b). We next tested the IN route, which is known to concentrate small, lipophilic compounds more efficiently in the brain than i.p. injections. This is likely because the olfactory and trigeminal nerve pathways allow IN-administered compounds to bypass the blood–brain barrier^45^, thus increasing compound bioavailability in the CNS noninvasively while reducing exposure to peripheral tissues. Since we were expecting improved efficacy, we opted for a lower IN dose of 3 mg/kg IS20. Using liquid chromatography/mass-spectrometry (LC/MS), our pharmacokinetic study found that the concentration of IN-injected IS20 reached 4 ng/mg tissue in the whole brain at 30 min before gradually decreasing to 1.5 ng/mg tissue at 90 min and to <0.8 ng/mg tissue at 6 h (Fig 4c), indicating that IN-injected IS20 was able to cross the blood–brain barrier and accumulate in the brain. Interestingly, the brain level of IS20 remained relatively high even 24 h post-IS20 treatment. ELISA detected concurrent increases in GDNF protein in both brain lysates and serum (Fig. 4d– e), with both responses peaking at 30 min post-IS20 treatment. Similar to i.p. injections of IS20, at 3 and 6 h post-IN administration of IS20, qPCR assays showed significant increases in GDNF mRNA expression in the SN and striatum (Fig. 4f). GDNF mediates pro-survival effects through its preferred receptor GFRα1, a glycosylphosphatidylinositol (GPI) protein, and its co-receptor RET, a receptor tyrosine kinase that transduces intracellular signals and whose decreased expression causes progressive degeneration of the nigrostriatal system^23,46^. Intriguingly, IS20 significantly increased the expression of GFRα1 and RET mRNAs in whole brain lysates (Fig. 4g– h). Collectively, these data show that both minimally invasive i.p. delivery and non-invasive IN delivery of IS20 upregulate GDNF and GDNF receptors in the mouse brain.

**Figure 4.**
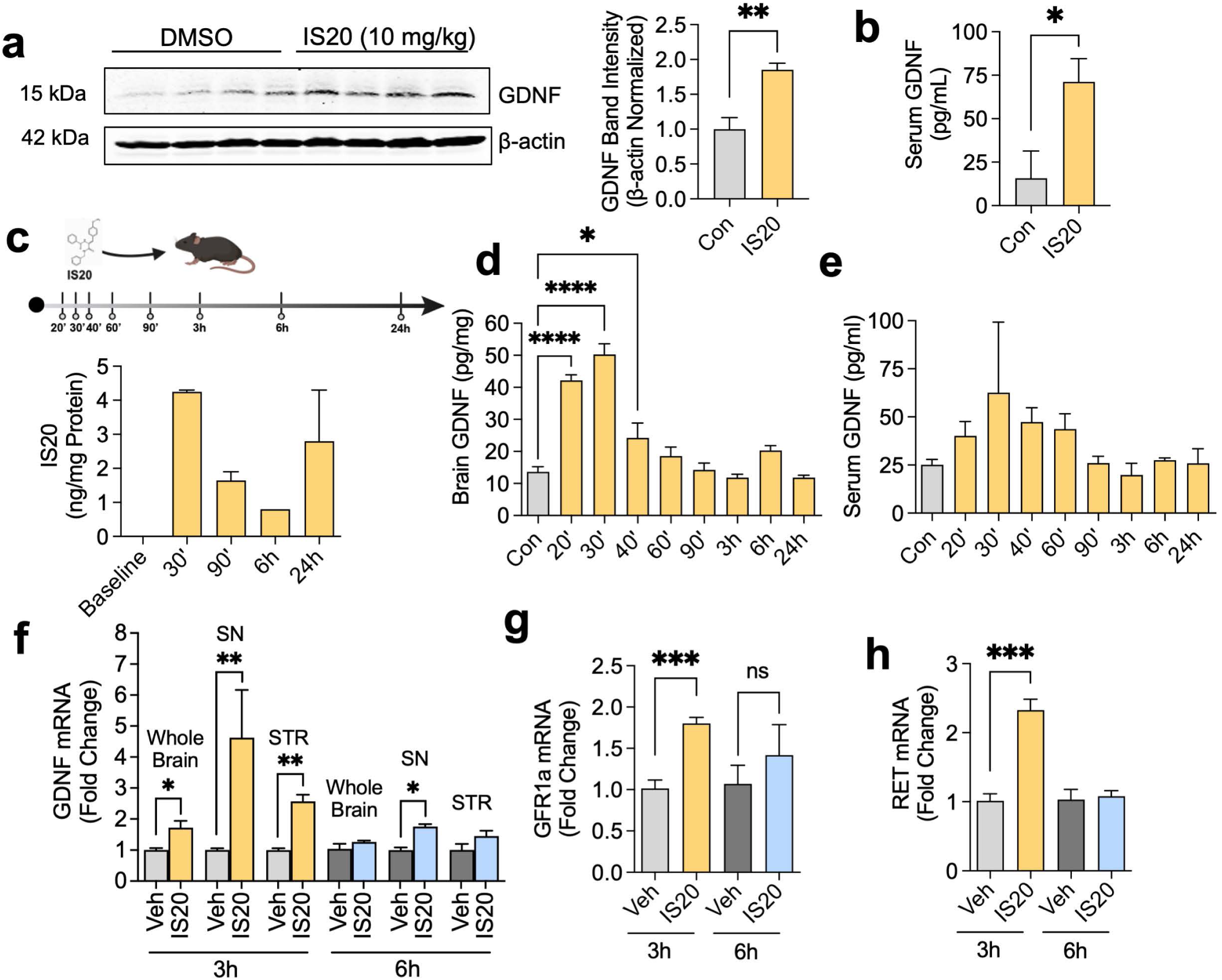
Intraperitoneal or intranasal administration of IS20 induces GDNF family neurotrophic factors in wild-type C57BL mice. (a) Western blot analysis probing for GDNF in substantia nigra (SN) lysates from mice intraperitoneally injected with 10 mg/kg IS20 8 h post-administration. (b) GDNF ELISA of blood serum 8 h after sacrifice of 10 mg/kg IS20-treated and vehicle-treated mice. (c) Mouse treatment paradigm (top) and liquid chromatography/mass spectrometry (LC/MS) analysis of IS20 in the brain of mice treated with IS20 (bottom). (d, e) ELISA assay of GDNF protein between 20 min to 24 h post intranasal IS20 (3 mg/kg) administration as measured in the (d) brain and (e) blood serum. (f) GDNF qPCR of the whole brain, SN, and striatum (STR) 3 h and 6 h after intranasal treatment of IS20 (3 mg/kg). (g, h) qPCR analysis of whole brain lysates collected 3 and 6 h post intranasal IS20 (3 mg/kg) administration, showing fold change in mRNA expression of (g) GFRa1 and (h) RET. *p≤0.05, **p<0.01, ***p<0.001.

### IS20 protects against 1-methyl-4-phenyl-1,2,3,6-tetrahydropyridine (MPTP)-induced DAergic cell death and preserves GDNF expression in C57BL/6 mice

Since intracranial delivery of exogenous rGDNF has shown effective neuroprotection^47^, we hypothesized that the early IS20-mediated induction of striatal GDNF would protect DAergic neurons against neurodegeneration in a mouse model of PD. To test this hypothesis, we co-injected C57BL/6 mice with the classic Parkinsonian toxicant MPTP and IS20 i.p. once daily for 5 d, with IS20 being injected 1 h after MPTP. After 5 d of the MPTP–IS20 co-injections, we injected only IS20 for another 7 d (Fig. 5a). This intervention strategy of combining co-treatments with post-MPTP treatments allowed us to examine IS20’s effect on the early neurodegenerative process. As confirmed by qPCR, MPTP treatment alone mildly reduced GDNF gene expression, while IS20 co-treatment significantly protected GDNF gene expression in MPTP-treated mice (Fig. 5b). In concordance with our above-mentioned *in vitro, ex vivo,* and *in vivo* pharmacodynamics experiments, the largest fold-change in GDNF gene upregulation occurred in mice treated with IS20 alone (Fig. 5b). HPLC analysis of striatal homogenates showed that IS20 treatment modestly but significantly attenuated the MPTP-induced depletion of DA and its metabolites HVA and DOPAC (Fig. 5c–e). Finally, to further determine the extent to which IS20-driven GDNF upregulation protected the nigral–striatal system against MPTP-induced lesioning, TH immunohistochemistry (IHC) and stereological counts of TH-positive DAergic neurons were performed on caudate-putamen and SN cryosections obtained from mice in each group. As expected, while MPTP-alone significantly reduced the number of TH-positive DAergic neurons in the SN, IS20 significantly preserved the TH-positive neuron count in MPTP/IS20-cotreated mice (Fig. 5f–g). TH IHC also showed that IS20 treatment significantly preserved DAergic nerve fibers damaged by MPTP treatments (Fig. 5g, representative sections from each group shown). These results suggest that IS20 treatment is protective against MPTP-induced neuronal and neurochemical deficits, possibly through enhanced GDNF signaling.

**Figure 5.**
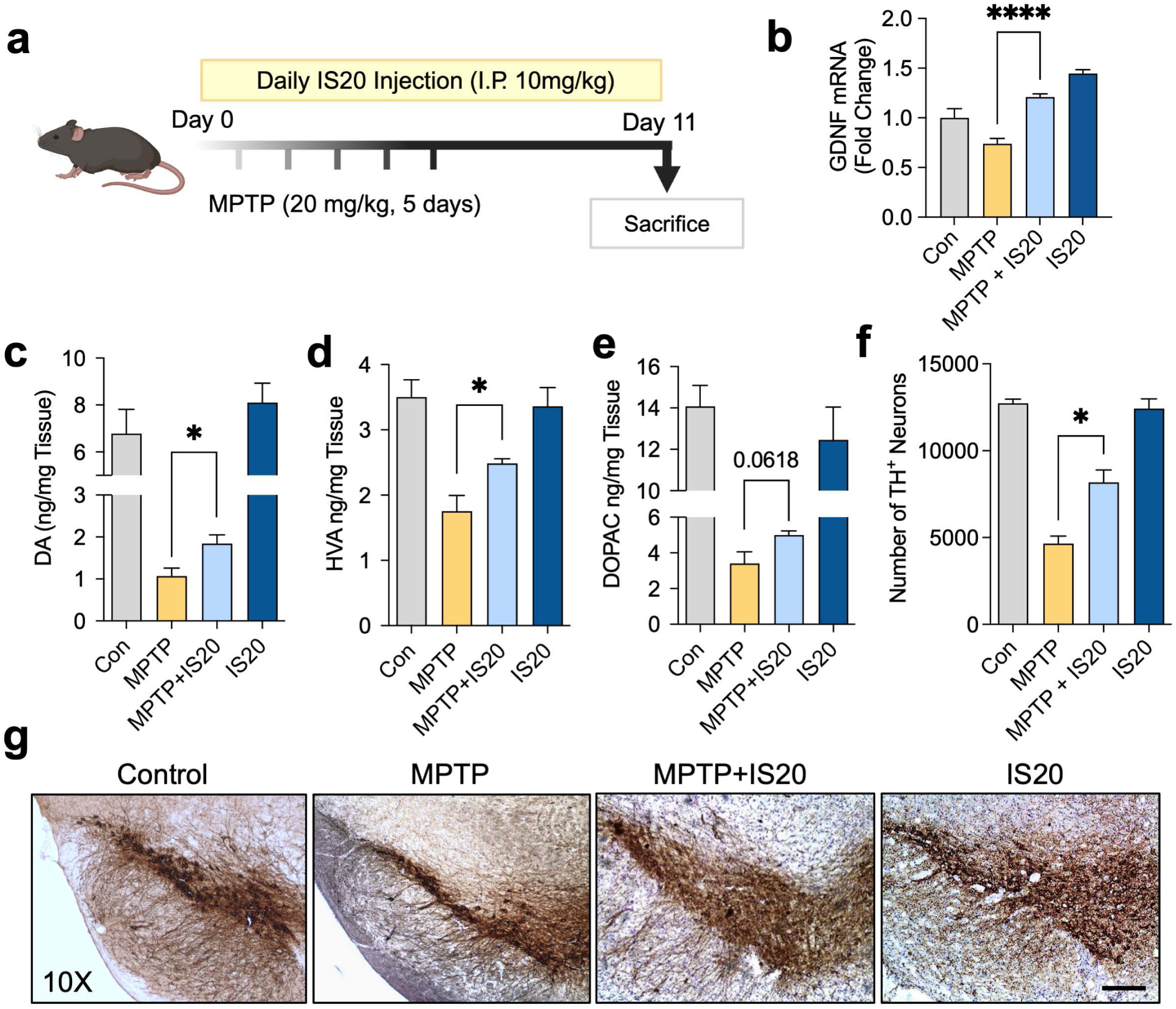
IS20 protects against MPTP-induced DAergic cell death in C57BL/6 mice and restores GDNF expression. (a) Treatment paradigm for C57BL/6 mice intraperitoneally injected with MPTP and IS20 treatments for 5 d and 12 d, respectively. (b) qPCR analysis showing GDNF gene expression of mice after 12 d of treatments with vehicle control, MPTP alone, MPTP-IS20 co-treatment, or IS20 alone. (c–e) HPLC analysis of striatal lysates from mice treated with vehicle control, MPTP alone, MPTP-IS20 co-treatment, or IS20 alone showing (c) DA, (d) homovanillic acid (HVA), and (e) DOPAC. (f) Software-assisted stereological count of TH+ DAergic neurons in the SNpc of IS20 co-treated mice compared to the MPTP group. (g) DAB immuno-staining images of TH+ neurons in the SNpc under 10x magnification (scale bar = 200 µm). Student’s t-tests were used to evaluate MPTP vs. MPTP+IS20. *p≤0.05, **p<0.01.

### IS20 protects against behavioral deficits and DAergic neurodegeneration in the MitoPark mouse model of PD

Because DAergic degeneration in the MPTP model stems from an acute neurotoxic insult and follows a rapid disease course, we next used the transgenic (Tg) MitoPark mouse model of PD, which more closely recapitulates PD’s chronically progressive neurodegenerative process. This genetic model of PD was rendered by cKO of mitochondrial transcription factor A (TFAM) driven by the Cre/LoxP system in DAergic neurons. The subsequent mitochondrial dysfunction results in PD-like symptoms, including a steady, progressive loss of DAergic neurons over several months, with the accompanying loss of motor capacity starting at around age 12–14 wk, as well as non-motor symptoms including olfactory deficits and depression. Littermates lacking the cKO of TFAM and DAT do not exhibit deficits and serve as ideal healthy controls. MitoPark mice age 13 wk were randomly chosen for IN treatment with vehicle or IS20 (3 mg/kg) for 4 wk (Fig. 6a).

**Figure 6.**
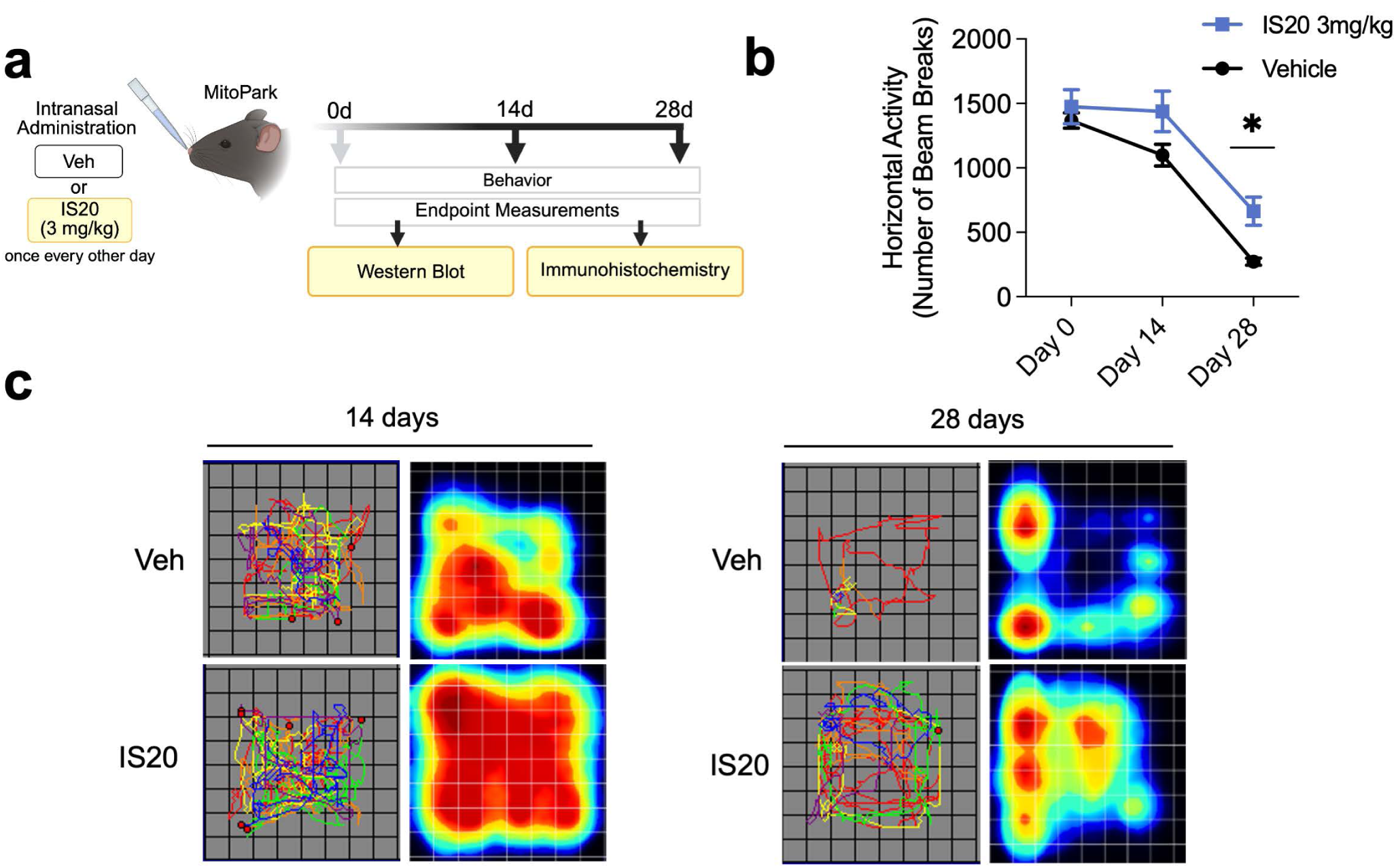
IS20 protects against behavioral deficits in MitoPark model. (a) Treatment paradigm for MitoPark mice treated with 3 mg/kg IS20 or vehicle via intranasal administration every other day for 28 d with biweekly behavioral monitoring using VersaMax. (b) Horizontal activity analysis of mice treated with IS20 (3 mg/kg) or vehicle after 14 and 28 d (n=2–3). (c) Representative tracing and heat plots of VersaMax analysis from 10-min activity monitoring of mice treated with IS20 (3 mg/kg) or vehicle after 14 and 28 d. *p≤0.05.

Using this model allowed us to test if a more chronic IN treatment regimen attenuates the onset and progression of behavioral and motor deficits in a more gradually progressive model. Starting at 15 wk (14 d into treatment), control MitoPark mice exhibited a drastic reduction in locomotor activity (Fig. 6b–c). However, MitoPark mice that received IN IS20 exhibited slightly more horizontal activity compared to vehicle controls, and by 28 d post-treatment, MitoPark mice that received IS20 performed significantly better (Fig. 6b–c).

The MitoPark mice were then sacrificed for tissues after treatment at age 17 wk. The SN of IS20-treated MitoPark mice showed significantly increased GNDF gene expression, as well as GDNF protein expression, as measured by qPCR and Western blot, respectively, when compared to vehicle-treated mice (Fig. 7a–b). TH protein levels in the MitoPark SN were also significantly protected by IS20 treatment, whereas PKR1 levels were unaffected (Fig. 7b–c). Since astrocyte-derived GDNF can reduce microglia-induced neuroinflammation^48,49^, we also assessed IBA1 protein levels and found a decrease in expression in the SN (Fig. 7b–c), potentially indicating a reduction in microglial activation in IS20-treated mice. To confirm the functional effects of IS20-induced GDNF upregulation on DAergic neurodegeneration in MitoPark mice, brain sections from 16-wk MitoPark mice were immunostained for TH to detect DAergic neurons. DAB immunostaining and stereological counts of TH-positive DAergic neurons revealed significantly higher neuronal counts in the SN when comparing IS20-to vehicle-treated MitoPark mice (Fig. 7d). Taken together, these experiments show that IS20 IN delivery could significantly protect against DAergic neuronal cell loss and behavioral deficits in the Tg MitoPark mouse model of PD.

**Figure 7.**
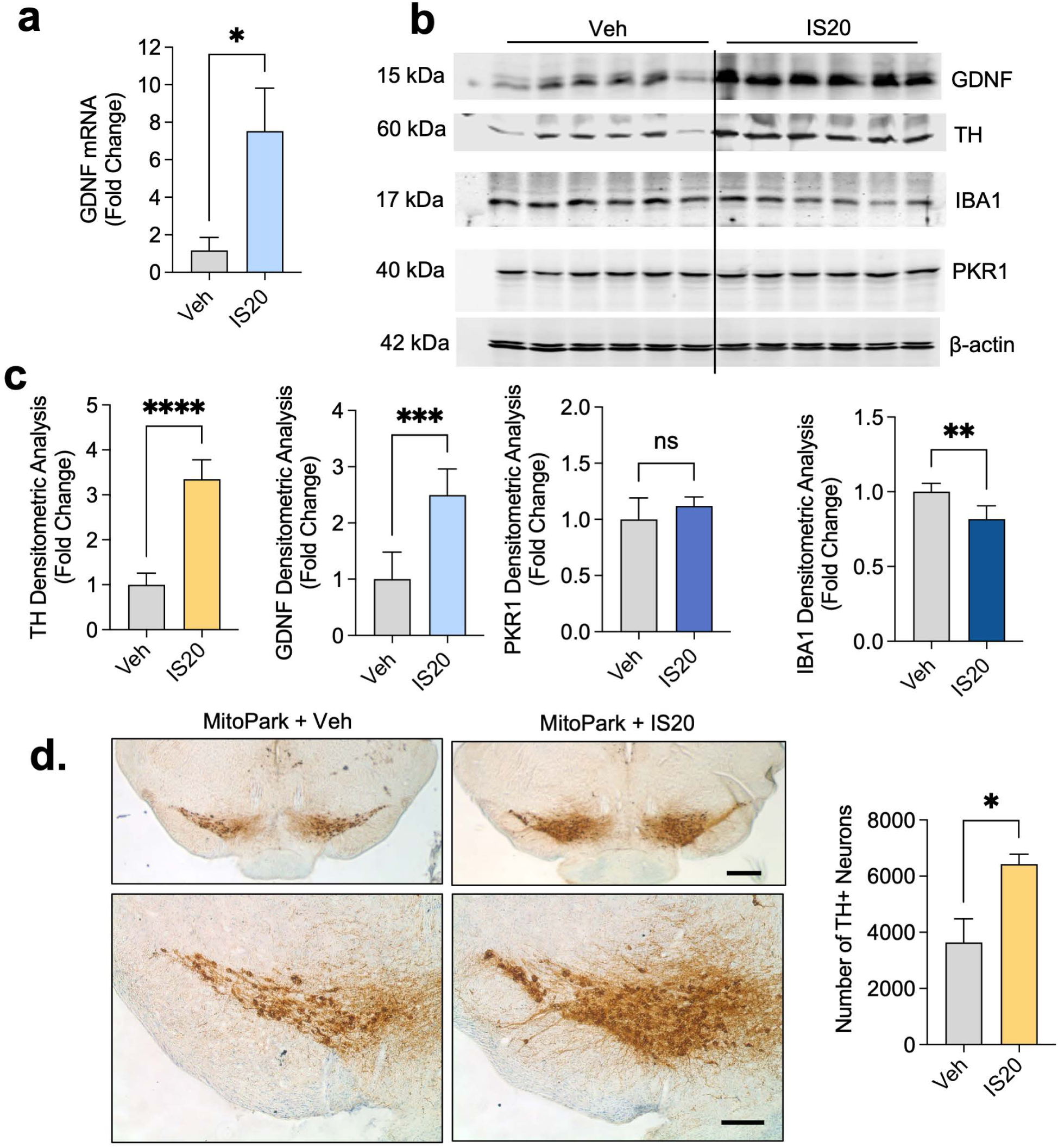
IS20 protects against DAergic neurodegeneration in MitoPark model. (a) qPCR of GDNF and of SN lysates from 13-wk-old MitoPark mice treated intranasally with IS20 (3 mg/kg) or vehicle control every other day for 28 d. (b) Western blot analysis probing for GDNF, TH, IBA1, and PKR1 in SN brain lysates from IS20-or vehicle-treated MitoPark mice; β-actin was used as loading control (n = 6). (c) Densitometric analysis of Western blots (shown in b) normalized to β-actin. (d) Tyrosine hydroxylase (TH) immunostaining of SN sections imaged at 2X (top) and 10X (bottom) magnifications with computer-assisted stereological counting of TH+ cells in the SNpc (right). *p≤0.05, **p<0.01, ***p<0.001 when compared to vehicle control group.

### IS20 activates AKT and p44/42 MAPK signaling pathways and induces pro-survival *Bcl2* gene expression

Knowing that AKT phosphorylation might induce GDNF upregulation, we explored IS20’s effects on cellular signal transduction in primary mouse astrocytes. IS20 rapidly induced both AKT and p44/42 phosphorylation (Fig. 8a) and promoted the expression of Bcl2 mRNA (Fig. 8b). Furthermore, in C57BL/6 mice 8 h after IP injection, IS20 induced AKT phosphorylation in the striatum (Fig. 8c). These experiments suggest mechanistically that IS20 promotes signaling transduction known to induce GDNF upregulation as well as other neuroprotective factors.

**Figure 8.**
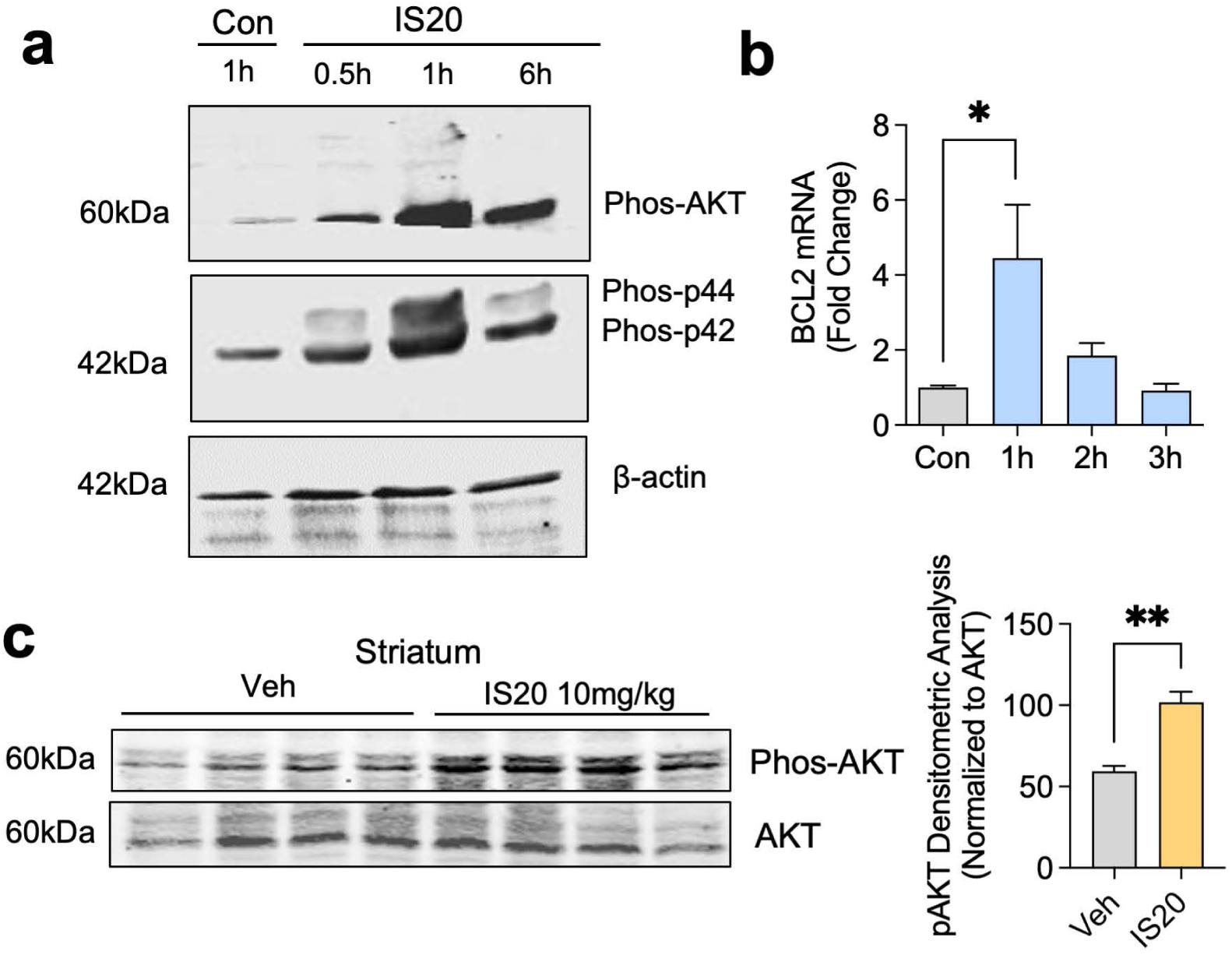
IS20 activates p44/42 and AKT pathways and induces pro-survival NRF2 and BCL2 gene expression. (a) Western blot analysis probing for phosphorylated AKT and phosphorylated p44/42 in primary mouse astrocytes treated with IS20 for 0.5–6 h. (b) qPCR of Bcl2 gene expression in primary mouse astrocytes treated with IS20. (c) Western blot analysis probing for phospho-AKT and AKT in striatum lysates from C57BL/6 mice at 8 h after IP injection of IS20. *p≤0.05, **p<0.01.

## Discussion

In this study, we demonstrated the potential for a small-molecule PKR1 receptor agonist to induce GDNF expression and to achieve neuroprotective effects in two rodent models of PD. In particular, we show that activation of PKR1 signaling through IS20 treatment quickly causes GDNF secretion and gene expression in primary astrocyte and organotypic slice cultures. In our experiments with C57BL/6 mice, IN-administered IS20 successfully crossed the blood–brain barrier, reached maximum concentration in around 30 min, and induced GDNF serum release and gene expression in the nigrostriatal system. Using a subacute MPTP mouse model of PD, we show that IS20 protects DAergic neurons against MPTP-induced neurodegeneration and helps to preserve striatal DA levels. Furthermore, we show in Tg MitoPark mice that a 4-wk course of IN-administered IS20 induced GDNF and TH levels in the SN, suppressed microglial activation, protected against DAergic neurodegeneration, and that these protective effects translate to functional improvements in locomotor activity for MitoPark mice. These findings suggest that pharmacological modulation of GDNF signaling is achievable with minimal side effects using IN administration of the PKR1 receptor agonist IS20.

PK2 activates PKR1 at an EC_50_ of 4.3 ± 1.3 nM and PKR2 at an EC_50_ of 7.3 ± 2.8 nM^50^. The apparently similar receptor affinities^51^ suggest that downstream effects induced by PK2 are determined by the expression levels of PKR1 and PKR2 on the cell surface^52^. We previously showed that PK2 signaling through PKR1 and PKR2 in neurons rapidly increases intracellular Ca^2+^ and induces phosphorylation of AKT and p44/42, leading to increases in BCL2 protein levels and mitochondrial biogenesis. We also developed an efficient AAV2/5-mediated gene delivery system, which produces robust, stable expression of PK2 in the striatum and protects SN neurons from MPTP-induced neurodegeneration. In astrocytes, activation of prokineticin signaling is likely modulated by the predominant expression of PKR1^36^. Our current study shows that the PKR1 receptor agonist IS20 rapidly increases intracellular Ca^2+^ and induces phosphorylation of AKT and p44/42 in astrocytes. AKT activation is a known pathway that results in secretion and upregulation of endogenous GDNF. Since both prokineticin and GDNF signaling depend on AKT and ERK signaling in DAergic neurons to mediate protective effects^35,46^, it is therefore possible that crosstalk between prokineticin and GDNF signaling mediates IS20’s pro-survival effects. Indeed, Tanabe *et al.* (2012)^53^ report that blocking AKT and p44/42 using API and PD98059, respectively, in PK2-treated neurons nullifies PK2’s protective effects, while blocking p44/42 using PD98059 in rat glioma cells prevents GDNF release. Their findings further suggest that prokineticin and GDNF signaling converge at AKT and p44/42 in DAergic neurons to promote neuronal survival.

Yamagata *et al.* (2002)^54^ revealed that GDNF is upregulated by ischemia and that astrocyte-derived GDNF protects neuronal cultures from cell death in cases of mitochondrial damage (see also C.-H. Lin et al., 2006). PK2 is also upregulated by hypoxia in primary cortical culture models of ischemia^55^; induces proliferation, survival and migration of capillary endothelial cells under hypoxic stress^56^; and supports mitochondrial biogenesis under MPP^+^-induced oxidative stress^35^. These previous findings support our hypothesis that PK2 is co-regulated in the brain with GDNF. Others also report significant crosstalk between PK2 and GDNF signaling in the development of the ENS^32^. Both PK2 and GDNF are also strong chemoattractants for projecting axons during neurogenesis and neuritogenesis. Additionally, PKR1 cKO leads to heart and kidney disorders due to deficits in angiogenesis, cell survival signaling, and mitochondrial and progenitor cell functions in these organs^57^. The involvement of both GDNF and PK2 signaling in PNS and CNS functions suggests crosstalk between their pathways.

Previous GDNF clinical trials concluded that although the potential for GDNF-based therapy remains high, two challenges need to be overcome: proper selection of animal-based PD models during preclinical testing and unwanted compensatory reactions or side effects associated with GDNF delivery^58^. MPTP has been the most frequently used Parkinsonian toxicant applied in the generation of animal models of PD^59,60^, ever since the accidental discovery decades ago of its ability to recapitulate the disease in humans^61,62^. Although subacute MPTP treatment (3–5 daily injections) in rodents produces PD-related neuropathology, this outcome does not progress gradually nor does it reliably reproduce the suite of behavioral deficits characterizing the human disease^63–65^. MPTP injections leave striatal DAergic neuron projections intact, which are then capable of retrogradely transporting GDNF to their somata in the SN. Yet, recent clinical data observed in post-mortem brains of PD patients 5 y post-diagnosis revealed that TH-positive DAergic innervations from the caudate–putamen completely disappeared, questioning the effectiveness of delivering GDNF only in the striatum of advanced PD patients, as employed by clinical trials conducted by Amgen^66–68^ and others^69,70^. Thus, the translational value of past studies relying only on acute neurodegeneration models that suggest targeting the striatum is both necessary and sufficient is confounded by their intact axonal transport machinery^71^. In contrast, MitoPark mice used in the current study lost striatal projections at age 17 wk, recapitulating conditions seen in advanced PD patients, thereby allowing critical assessments of preclinical functional endpoints. This current understanding sheds light on earlier studies reporting that GDNF administration by intranigral injection could mitigate neuronal cell loss, but did not protect against loss of terminal projections^56,72^. And by extension, intrastriatal injection of AAV-GDNF could protect against degeneration of both neuronal cell bodies and projections if administered before neuronal cell loss^73^. This is presumably because GDNF must be retrogradely transported to the neurosomata for RET signaling to mediate pro-survival effects^23^. Therefore, high GDNF expression in the striatum is less effective if GDNF’s retrograde transport is disrupted due to significantly degenerated DAergic neuronal projections. In this case, increasing GDNF levels in both the striatum and SN might prove to be more effective in promoting GDNF-induced pro-survival effects^74^. Indeed, success has been achieved using AAV2-mediated neurotrophic factor gene transfer in both the striatum and nigra^73,75^. In our current study, IS20-induced GDNF expression levels were upregulated in both the striatum and SN of MitoPark mice, allowing astrocyte-secreted GDNF to protect both DAergic projections in the striatum as well as neurosomata in the SN. Significantly more striatal projections are preserved in MitoPark mice treated with IS20. Together, these data indicate that performing preclinical studies in animal models that recapitulate the gradual progression of PD pathology could improve the transition from translational research to clinical trials for PD and reduce the attrition of candidate therapies.

Most current GDNF gene delivery trials use AAV2, which has low immunogenicity and has proven safe for use in humans, but transduces predominantly neurons^76,77^. However, in the injured brain, GDNF is mainly produced by astrocytes, not by neurons^78,79^, as when astrocytes secrete GDNF during ischemia as a compensatory response^80^. Astrocytes also provide the major source of GDNF in the SN of PD patients’ midbrains^28,29^, where astrocytes and microglia outnumber neurons 10 to 1^49,81^. Early studies isolated nigral GDNF from Type 1 astrocytes and found that astrocyte-derived GDNF enhanced DAergic neuron survival when nigral astrocytes were co-cultured with DAergic neurons as support cell monolayers^11^. Astrocytes are also known to secrete other neurotrophic factors, such as CDNF, BDNF, and NGF. Moreover, in the gut, enteric glia cells are the major source of GDNF^82,83^. Therefore, neuroprotective efforts targeting only neurons for long-term neuronal viability are unlikely to succeed if supportive astrocytes do not provide proper neurotrophic and metabolic enrichments^84^. Several studies have found that intrastriatal, viral-mediated GDNF gene delivery causes aberrant neurite growth towards the site of application^23,39^, suggesting that mimicking the endogenous mode of expression is important to achieve functional effects. Drinkut et al. (2012)^28^ focused on viral delivery of GDNF into astrocytes, achieving localized, yet satisfactory neuroprotection. Building upon these studies, we found here that targeted GDNF upregulation in astrocytes could be achieved pharmacologically using a non-invasive route of administration, suggesting that additional research efforts should be redirected back to astrocytes.

The neuroprotective effects afforded by IS20 could also be mediated through the reduction of microglial IBA1 expression in the SN, which has a high density of microglia^85^, whose activation is affected by PD pathology^86^. But here, too, the role of astrocytic GDNF appears to extend to neuroinflammation associated with neurodegeneration as it can potently inhibit excessive production of reactive oxygen species from microglia in zymosan A-stimulated midbrain microglia cultures^49^ and LPS-stimulated primary midbrain neuron–glia cultures^87^. Thus, astrocyte-derived GDNF could mediate protective effects by its anti-inflammatory effects on activated microglia^86^. In line with previous studies, we show that IS20 treatment reduced IBA1 expression in the SN of 17-wk-old MitoPark mice. However, constitutive GDNF expression from viral transgene expression can downregulate TH expression^24^, possibly due to feedback loops between GDNF and DA reuptake^88,89^. To counteract such unwanted compensatory effects, a discontinuous GDNF delivery paradigm could allow GDNF to return to basal levels^28,58^. Our pharmacodynamic studies in healthy control mice found that IS20-induced GDNF expression in the striatum and SN, as well as the whole brain, returned to basal levels 6 h post-treatment, thereby avoiding constitutive GDNF expression. Consequently, we saw increases in TH in MitoPark mice treated with IS20. Yet another possible cause of an unwanted compensatory effect, driven by viral transgenes in other studies^26^, results from GDNF expression that is at least one or two orders of magnitude higher, In our study, we have seen 2-fold and 3-fold increases in GDNF expression induced by IS20 in the striatum and SN, respectively, and a 0.5-fold increase in GDNF expression in whole brain relative to control levels. Furthermore, we did not see overt signs of toxicity or a significant reduction in body weight with the administered doses, routes, and study duration. This is in agreement with a previously published study, where subchronic i.p. administrations of PK2 protein, which activates PKR1 and PKR2 in a relatively non-selective fashion^90^, were administered systemically in C57BL/6 mice without causing serious toxicity^91^.

In summary, IS20 administration could pharmacologically upregulate GDNF levels in the nigrostriatal pathway, preserving DAergic neurons and functional innervations in neuroprotective paradigms applied in PD animal models. Importantly, the relative safety profile of IS20 makes it a promising pharmacological candidate for PD therapy.

## Materials and Methods

### Calcium Assays

Intracellular calcium mobilization was assessed using the Fluo-4 NW Calcium Assay Kit (F36206, Thermo Fisher, Waltham, MA) according to manufacturer’s instructions. Briefly, CHO cells were grown on 96-well plates the previous day. FLIPR (Molecular Devices, San Jose, CA) was used to inject dissolved IS20 into the cell plate wells, which were read on a FlexStation multi-mode microplate reader (Molecular Devices).

### Animal handling

All animal procedures were approved by Iowa State University’s Institutional Animal Care and Use Committee (IACUC). All mice were housed under a 12-h light cycle in a climate-controlled mouse facility (22±1 °C) with food and water available *ad libitum*. Male C57BL/6 mice were pre-screened during behavior assessments for normal baseline performance before being randomly assigned to experimental groups. None of the animals manifested wounds or any abnormal behaviors that would have warranted their removal from the study. Investigators involved with data collection and analysis were not blinded to group allocation.

### MPTP injections

Over a 5-day period, one i.p. injection of 18 mg/kg MPTP was administered every day to each mouse in the MPTP group, and an equal volume of saline (vehicle) was similarly injected into each mouse in the control group.

### Behavior monitoring

All groups were monitored for voluntary, exploratory locomotor activity using the automated VersaMax system and (analyzer, model VMAUSB, AccuScan, Columbus, OH) connected to VersaMax motion-detection hardware (monitor, model RXYZCM-16). The 40×40-cm square monitor was partitioned into four, 20×20-cm square open-field arenas. The VersaMax system is capable of tracking two animals simultaneously when using two diagonally opposed arenas that did not share a wall of infra-red (IR) beams. Each animal was allowed to acclimate to its arena for 2 min, after which its spontaneous locomotor activities (horizontal activity, vertical activity, and speed), monitored as IR beam-breaks, were recorded for 10 min. Fusion software was used to generate heat maps (Omnitech Electronics, Columbus, OH).

### MitoPark mouse model of PD

MitoPark mice were a kind gift from Dr Nils-Göran Larsson of the Karolinska Institute, Stockholm, Sweden, from his laboratory at the Max Planck Institute for Biology of Ageing. The Tg MitoPark mouse model is created by TFAM inactivation specifically in DA neurons by cKO through control of the DA transporter promoter. The mice used were from our MitoPark breeding colony at Iowa State University. After behavioral experiments were performed at the predetermined ages, mice were euthanized at the predetermined endpoints via CO_2_ asphyxiation, as outlined in approved IACUC protocols, and samples processed for either RT-qPCR or Western blotting.

### Western blot

Dissected brain regions were collected in Eppendorf tubes and flash frozen. To isolate total protein, RIPA buffer containing sodium orthovanadate and Protease and Phosphatase Inhibitor Cocktail (Thermo Fisher 78440) was added to each tube and homogenized using a tissue homogenizer. Dissolved total lysate was centrifuged at 12,000 x g for 60 min to remove cellular debris. Normalized protein samples were loaded into each well of a 12-well plate and were separated by sodium dodecyl sulfate (SDS) gel electrophoresis (100 V, 90 min), using Any kD Mini-PROTEAN resolving gel (4569035, Bio-Rad, Hercules, CA). Proteins on SDS gels were then transferred to a nitrocellulose membrane (26 V, overnight) and blocked for 1 h using fluorescent Western blocking buffer (Rockland Immunochemicals, Pottstown, PA). Primary antibodies diluted in blocking buffer with 0.05% Tween 20 were then added to the membranes and incubated overnight at 4 °C. Next day, primary antibodies were removed, and blots were washed (7 times, 5 min each) in PBS wash buffer containing 0.05% Tween 20 (PBST). Secondary antibody (infrared dye-tagged) was added for 1 h. Blots were then triple-washed in PBST and once in PBS. The protein β-actin was used as a loading control. Membranes were scanned using the Odyssey IR imaging system (LI-COR, Lincoln, NE) and digital images were captured via a LI-COR Odyssey imager. Densitometric analysis was done using ImageJ software.

### IHC

Cells were fixed in 4% paraformaldehyde, blocked with BSA and Triton-X to permeabilize the cell membranes. Primary antibodies were diluted as follows: rabbit anti-TH (1:1600), rabbit anti-PK2 (1:500), goat anti-GDNF (1:500), goat anti-IBA1 (1:1000), and rabbit anti-GFAP (1:1000). Alexa Fluor fluorescent secondary antibodies (Thermo Fisher) were used against primary antibodies for fluorescent imaging. For DAB staining, HRP-conjugated secondary antibodies were used against primary antibodies, and VECTASTAIN Elite ABC HRP Kit (Vector Labs, Newark, CA) was used for conjugating HRP to secondary antibodies. DAB was used for colorimetric detection.

### SYBR Green RT-qPCR

To obtain total RNA, tissue lysis buffer from the Absolutely RNA Miniprep kit (Agilent Technologies) was added to dissected, region-specific brain tissue samples together with beta-mercaptoethanol as a reducing agent to preserve RNA. A tissue homogenizer was used to dissolve tissue into the lysis buffer, and lysates were processed according to kit manufacturer’s instructions. Total RNA isolated from each sample was quantified using NanoDrop (Thermo Scientific) to determine RNA concentration and purity. First, strand cDNA synthesis was performed using the Affinity Script qPCR cDNA synthesis system (Agilent Technologies) with 1 µg of total RNA in the reaction mixture. Real-time PCR was performed with the RT^2^ SYBR Green master mix (Qiagen, Germantown, MD) using diluted cDNA and QuantiTect (Qiagen) mouse primer sets. The 18S rRNA gene (mouse) was used as the housekeeping gene to normalize each sample. The amount of each template was optimized empirically to maximize efficiency without inhibiting the PCR reaction. After the last cycle of the qPCR run, dissociation curves were run to ensure a single amplicon peak was obtained, indicating primer specificity. The results are reported as fold change in gene expression derived with the ΔΔC_t_ method, using the threshold cycle (C_t_) value for the housekeeping gene and for the respective gene of interest in each sample. Control animals serve as the baseline for fold change.

### Lentivirus production

The PK2 lentivirus expression vector (OriGene, Rockville, MD) was mixed with the MISSION Lentiviral Packaging Mix (SHP001, Sigma, St. Louis, MO) according to manufacturer’s protocols, and co-transfected into 293FT cells to package the virus. After 24 h post-transfection, the supernatant, which contains the virus, was collected. The second harvesting of virus was done 48 h post-transfection. The Lenti-X™ p24 Rapid Titer Kit (632200, Clontech, Mountain View, CA) was used to titer the lentivirus.

### HPLC analysis of striatal DA levels

HPLC samples were processed as previously described^35^ (Gordon et al., 2016). Briefly, mice were euthanized, striata were collected and neurotransmitters were extracted in 0.2 M perchloric acid solution containing 0.05% Na_2_EDTA, 0.1% Na_2_S_2_O_5_ and isoproterenol (internal standard). DA and metabolites were separated isocratically by a reversed-phase column with a flow rate of 0.6 ml min^−1^ using a Dionex Ultimate 3000 HPLC system (pump model ISO-3100SD, Thermo Scientific) equipped with a refrigerated automatic sampler (model WPS-3000TSL). The CoulArray (model 5600A) electrochemical detection system was coupled with an analytical cell (microdialysis cell model 5014B) and a guard cell (model 5020). Data acquisition and analysis were performed using Chromeleon 7 and ESA CoulArray 3.10 HPLC Software.

## Data analysis

All *in vitro* data were determined from at least two biologically independent experiments, each done with a minimum of three biological replicates. Data analysis used Prism 10.5.0 software (GraphPad Software, San Diego, CA) to perform one-way ANOVA and then Bonferroni’s post-tests or two-way ANOVA to compare all treatment groups. Student’s t-test (two-tailed) was used when two groups were being compared. Differences with p≤0.05 were considered statistically significant.

## Acknowledgements

This work was supported by National Institutes of Health grant R01 ES034196. Other sources include the Johnny Isakson Endowment and the Coach Mark Richt Neurological Disease Research Fund to A.G.K. and A.K.

## References

1 Barzilai, A. & Melamed, E. Molecular mechanisms of selective dopaminergic neuronal death in Parkinson’s disease. Trends Mol Med 9, 126–132 (2003). 10.1016/s1471-4914(03)00020-0

2 Jarraya, B. et al. Dopamine gene therapy for Parkinson’s disease in a nonhuman primate without associated dyskinesia. Sci Transl Med 1, 2ra4 (2009). 10.1126/scitranslmed.3000130

3 Meissner, W. G. et al. Priorities in Parkinson’s disease research. Nat Rev Drug Discov 10, 377–393 (2011). 10.1038/nrd3430

4 Birkmayer, W. & Hornykiewicz, O. The L-3,4-dioxyphenylalanine (DOPA)-effect in Parkinson-akinesia. Wien Klin Wochenschr 73, 787–788 (1961).

5 Won, S. Y. et al. Nigral dopaminergic PAK4 prevents neurodegeneration in rat models of Parkinson’s disease. Sci Transl Med 8, 367ra170 (2016). 10.1126/scitranslmed.aaf1629

6 Rascol, O. et al. A five-year study of the incidence of dyskinesia in patients with early Parkinson’s disease who were treated with ropinirole or levodopa. N Engl J Med 342, 1484–1491 (2000). 10.1056/NEJM200005183422004

7 Enomoto, H. et al. GFR alpha1-deficient mice have deficits in the enteric nervous system and kidneys. Neuron 21, 317–324 (1998). 10.1016/s0896-6273(00)80541-3

8 Young, H. M. et al. GDNF is a chemoattractant for enteric neural cells. Dev Biol 229, 503–516 (2001). 10.1006/dbio.2000.0100

9 Uesaka, T., Nagashimada, M. & Enomoto, H. GDNF signaling levels control migration and neuronal differentiation of enteric ganglion precursors. J Neurosci 33, 16372–16382 (2013). 10.1523/JNEUROSCI.2079-13.2013

10 Pascual, A. et al. Absolute requirement of GDNF for adult catecholaminergic neuron survival. Nat Neurosci 11, 755–761 (2008). 10.1038/nn.2136

11 O’Malley, E. K., Sieber, B. A., Black, I. B. & Dreyfus, C. F. Mesencephalic type I astrocytes mediate the survival of substantia nigra dopaminergic neurons in culture. Brain Res 582, 65–70 (1992). 10.1016/0006-8993(92)90317-3

12 Lin, L. F., Doherty, D. H., Lile, J. D., Bektesh, S. & Collins, F. GDNF: a glial cell line-derived neurotrophic factor for midbrain dopaminergic neurons. Science 260, 1130–1132 (1993). 10.1126/science.8493557

13 Beck, K. D. et al. Mesencephalic dopaminergic neurons protected by GDNF from axotomy-induced degeneration in the adult brain. Nature 373, 339–341 (1995). 10.1038/373339a0

14 Bilang-Bleuel, A. et al. Intrastriatal injection of an adenoviral vector expressing glial-cell-line-derived neurotrophic factor prevents dopaminergic neuron degeneration and behavioral impairment in a rat model of Parkinson disease. Proc Natl Acad Sci U S A 94, 8818–8823 (1997). 10.1073/pnas.94.16.8818

15 Bjorklund, A., Rosenblad, C., Winkler, C. & Kirik, D. Studies on neuroprotective and regenerative effects of GDNF in a partial lesion model of Parkinson’s disease. Neurobiol Dis 4, 186–200 (1997). 10.1006/nbdi.1997.0151

16 Kordower, J. H. et al. Neurodegeneration prevented by lentiviral vector delivery of GDNF in primate models of Parkinson’s disease. Science 290, 767–773 (2000). 10.1126/science.290.5492.767

17 Tseng, J. L., Baetge, E. E., Zurn, A. D. & Aebischer, P. GDNF reduces drug-induced rotational behavior after medial forebrain bundle transection by a mechanism not involving striatal dopamine. J Neurosci 17, 325–333 (1997). 10.1523/JNEUROSCI.17-01-00325.1997

18 Domanskyi, A., Saarma, M. & Airavaara, M. Prospects of Neurotrophic Factors for Parkinson’s Disease: Comparison of Protein and Gene Therapy. Hum Gene Ther 26, 550–559 (2015). 10.1089/hum.2015.065

19 Gill, S. S. et al. Direct brain infusion of glial cell line-derived neurotrophic factor in Parkinson disease. Nat Med 9, 589–595 (2003). 10.1038/nm850

20 Hovland, D. N., Jr., et al. Six-month continuous intraputamenal infusion toxicity study of recombinant methionyl human glial cell line-derived neurotrophic factor (r-metHuGDNF in rhesus monkeys. Toxicol Pathol 35, 1013–1029 (2007). 10.1177/01926230701481899

21 Blits, B. & Petry, H. Perspective on the Road toward Gene Therapy for Parkinson’s Disease. Front Neuroanat 10, 128 (2016). 10.3389/fnana.2016.00128

22 Remy, P. Biotherapies for Parkinson disease. Rev Neurol (Paris) 170, 763–769 (2014). 10.1016/j.neurol.2014.10.002

23 Tenenbaum, L. & Humbert-Claude, M. Glial Cell Line-Derived Neurotrophic Factor Gene Delivery in Parkinson’s Disease: A Delicate Balance between Neuroprotection, Trophic Effects, and Unwanted Compensatory Mechanisms. Front Neuroanat 11, 29 (2017). 10.3389/fnana.2017.00029

24 Tereshchenko, J., Maddalena, A., Bahr, M. & Kugler, S. Pharmacologically controlled, discontinuous GDNF gene therapy restores motor function in a rat model of Parkinson’s disease. Neurobiol Dis 65, 35–42 (2014). 10.1016/j.nbd.2014.01.009

25 Caumont, A. S., Octave, J. N. & Hermans, E. Specific regulation of rat glial cell line-derived neurotrophic factor gene expression by riluzole in C6 glioma cells. J Neurochem 97, 128–139 (2006). 10.1111/j.1471-4159.2006.03711.x

26 Ibanez, C. F. & Andressoo, J. O. Biology of GDNF and its receptors - Relevance for disorders of the central nervous system. Neurobiol Dis 97, 80–89 (2017). 10.1016/j.nbd.2016.01.021

27 Dougherty, K. D., Dreyfus, C. F. & Black, I. B. Brain-derived neurotrophic factor in astrocytes, oligodendrocytes, and microglia/macrophages after spinal cord injury. Neurobiol Dis 7, 574–585 (2000). 10.1006/nbdi.2000.0318

28 Drinkut, A., Tereshchenko, Y., Schulz, J. B., Bahr, M. & Kugler, S. Efficient gene therapy for Parkinson’s disease using astrocytes as hosts for localized neurotrophic factor delivery. Mol Ther 20, 534–543 (2012). 10.1038/mt.2011.249

29 Mogi, M. et al. Glial cell line-derived neurotrophic factor in the substantia nigra from control and parkinsonian brains. Neurosci Lett 300, 179–181 (2001). 10.1016/s0304-3940(01)01577-4

30 Laustsen, A. H., Lomonte, B., Lohse, B., Fernandez, J. & Gutierrez, J. M. Unveiling the nature of black mamba (Dendroaspis polylepis) venom through venomics and antivenom immunoprofiling: Identification of key toxin targets for antivenom development. J Proteomics 119, 126–142 (2015). 10.1016/j.jprot.2015.02.002

31 Ngan, E. S. et al. Prokineticin-1 modulates proliferation and differentiation of enteric neural crest cells. Biochim Biophys Acta 1773, 536–545 (2007). 10.1016/j.bbamcr.2007.01.013

32 Ngan, E. S. et al. Prokineticin-1 (Prok-1) works coordinately with glial cell line-derived neurotrophic factor (GDNF) to mediate proliferation and differentiation of enteric neural crest cells. Biochim Biophys Acta 1783, 467–478 (2008). 10.1016/j.bbamcr.2007.09.005

33 Ruiz-Ferrer, M. et al. Expression of PROKR1 and PROKR2 in human enteric neural precursor cells and identification of sequence variants suggest a role in HSCR. PLoS One 6, e23475 (2011). 10.1371/journal.pone.0023475

34 Ng, K. L. et al. Dependence of olfactory bulb neurogenesis on prokineticin 2 signaling. Science 308, 1923–1927 (2005). 10.1126/science.1112103

35 Gordon, R. et al. Prokineticin-2 upregulation during neuronal injury mediates a compensatory protective response against dopaminergic neuronal degeneration. Nat Commun 7, 12932 (2016). 10.1038/ncomms12932

36 Koyama, Y. et al. Expression of prokineticin receptors in mouse cultured astrocytes and involvement in cell proliferation. Brain Res 1112, 65–69 (2006). 10.1016/j.brainres.2006.07.013

37 Gasser, A. et al. Discovery and cardioprotective effects of the first non-Peptide agonists of the G protein-coupled prokineticin receptor-1. PLoS One 10, e0121027 (2015). 10.1371/journal.pone.0121027

38 Georgievska, B., Kirik, D. & Bjorklund, A. Overexpression of glial cell line-derived neurotrophic factor using a lentiviral vector induces time-and dose-dependent downregulation of tyrosine hydroxylase in the intact nigrostriatal dopamine system. J Neurosci 24, 6437–6445 (2004). 10.1523/JNEUROSCI.1122-04.2004

39 Georgievska, B., Kirik, D. & Bjorklund, A. Aberrant sprouting and downregulation of tyrosine hydroxylase in lesioned nigrostriatal dopamine neurons induced by long-lasting overexpression of glial cell line derived neurotrophic factor in the striatum by lentiviral gene transfer. Exp Neurol 177, 461–474 (2002). 10.1006/exnr.2002.8006

40 Sajadi, A., Bauer, M., Thony, B. & Aebischer, P. Long-term glial cell line-derived neurotrophic factor overexpression in the intact nigrostriatal system in rats leads to a decrease of dopamine and increase of tetrahydrobiopterin production. J Neurochem 93, 1482–1486 (2005). 10.1111/j.1471-4159.2005.03139.x

41 Chauhan, N. B., Siegel, G. J. & Lee, J. M. Depletion of glial cell line-derived neurotrophic factor in substantia nigra neurons of Parkinson’s disease brain. J Chem Neuroanat 21, 277–288 (2001). 10.1016/s0891-0618(01)00115-6

42 Hunot, S. et al. Glial cell line-derived neurotrophic factor (GDNF) gene expression in the human brain: a post mortem in situ hybridization study with special reference to Parkinson’s disease. J Neural Transm (Vienna) 103, 1043–1052 (1996). 10.1007/BF01291789

43 Gasser, A. et al. Prokineticin Receptor-1 Signaling Inhibits Dose-and Time-Dependent Anthracycline-Induced Cardiovascular Toxicity Via Myocardial and Vascular Protection. JACC CardioOncol 1, 84–102 (2019). 10.1016/j.jaccao.2019.06.003

44 Ngan, E. S. & Tam, P. K. Prokineticin-signaling pathway. Int J Biochem Cell Biol 40, 1679–1684 (2008). 10.1016/j.biocel.2008.03.010

45 Hernando, S. et al. Intranasal Administration of TAT-Conjugated Lipid Nanocarriers Loading GDNF for Parkinson’s Disease. Mol Neurobiol 55, 145–155 (2018). 10.1007/s12035-017-0728-7

46 Sariola, H. & Saarma, M. Novel functions and signalling pathways for GDNF. J Cell Sci 116, 3855–3862 (2003). 10.1242/jcs.00786

47 Wang, Y., Lin, S. Z., Chiou, A. L., Williams, L. R. & Hoffer, B. J. Glial cell line-derived neurotrophic factor protects against ischemia-induced injury in the cerebral cortex. J Neurosci 17, 4341–4348 (1997). 10.1523/JNEUROSCI.17-11-04341.1997

48 Chen, S. H. et al. Microglial regulation of immunological and neuroprotective functions of astroglia. Glia 63, 118–131 (2015). 10.1002/glia.22738

49 Rocha, S. M., Cristovao, A. C., Campos, F. L., Fonseca, C. P. & Baltazar, G. Astrocyte-derived GDNF is a potent inhibitor of microglial activation. Neurobiol Dis 47, 407–415 (2012). 10.1016/j.nbd.2012.04.014

50 Lin, D. C. et al. Identification and molecular characterization of two closely related G protein-coupled receptors activated by prokineticins/endocrine gland vascular endothelial growth factor. J Biol Chem 277, 19276–19280 (2002). 10.1074/jbc.M202139200

51 Zhou, Q. Y. The prokineticins: a novel pair of regulatory peptides. Mol Interv 6, 330–338 (2006). 10.1124/mi.6.6.6

52 Attramadal, H. Prokineticins and the heart: diverging actions elicited by signalling through prokineticin receptor-1 or-2. Cardiovasc Res 81, 3–4 (2009). 10.1093/cvr/cvn306

53 Tanabe, K., Matsushima-Nishiwaki, R., Iida, M., Kozawa, O. & Iida, H. Involvement of phosphatidylinositol 3-kinase/Akt on basic fibroblast growth factor-induced glial cell line-derived neurotrophic factor release from rat glioma cells. Brain Res 1463, 21–29 (2012). 10.1016/j.brainres.2012.04.057

54 Yamagata, K. et al. Differential regulation of glial cell line-derived neurotrophic factor (GDNF) mRNA expression during hypoxia and reoxygenation in astrocytes isolated from stroke-prone spontaneously hypertensive rats. Glia 37, 1–7 (2002). 10.1002/glia.10003

55 Cheng, M. Y. et al. Prokineticin 2 is an endangering mediator of cerebral ischemic injury. Proc Natl Acad Sci U S A 109, 5475–5480 (2012). 10.1073/pnas.1113363109

56 LeCouter, J. et al. The endocrine-gland-derived VEGF homologue Bv8 promotes angiogenesis in the testis: Localization of Bv8 receptors to endothelial cells. Proc Natl Acad Sci U S A 100, 2685–2690 (2003). 10.1073/pnas.0337667100

57 Boulberdaa, M. et al. Genetic inactivation of prokineticin receptor-1 leads to heart and kidney disorders. Arterioscler Thromb Vasc Biol 31, 842–850 (2011). 10.1161/ATVBAHA.110.222323

58 Taylor, H. et al. Clearance and toxicity of recombinant methionyl human glial cell line-derived neurotrophic factor (r-metHu GDNF) following acute convection-enhanced delivery into the striatum. PLoS One 8, e56186 (2013). 10.1371/journal.pone.0056186

59 Beal, M. F. Experimental models of Parkinson’s disease. Nat Rev Neurosci 2, 325–334 (2001). 10.1038/35072550

60 Przedborski, S. et al. The parkinsonian toxin 1-methyl-4-phenyl-1,2,3,6-tetrahydropyridine (MPTP): a technical review of its utility and safety. J Neurochem 76, 1265–1274 (2001). 10.1046/j.1471-4159.2001.00183.x

61 Davis, G. C. et al. Chronic Parkinsonism secondary to intravenous injection of meperidine analogues. Psychiatry Res 1, 249–254 (1979). 10.1016/0165-1781(79)90006-4

62 Langston, J. W., Ballard, P., Tetrud, J. W. & Irwin, I. Chronic Parkinsonism in humans due to a product of meperidine-analog synthesis. Science 219, 979–980 (1983). 10.1126/science.6823561

63 Rommelfanger, K. S. et al. Norepinephrine loss produces more profound motor deficits than MPTP treatment in mice. Proc Natl Acad Sci U S A 104, 13804–13809 (2007). 10.1073/pnas.0702753104

64 Schober, A. Classic toxin-induced animal models of Parkinson’s disease: 6-OHDA and MPTP. Cell Tissue Res 318, 215–224 (2004). 10.1007/s00441-004-0938-y

65 Tillerson, J. L., Caudle, W. M., Reveron, M. E. & Miller, G. W. Detection of behavioral impairments correlated to neurochemical deficits in mice treated with moderate doses of 1-methyl-4-phenyl-1,2,3,6-tetrahydropyridine. Exp Neurol 178, 80–90 (2002). 10.1006/exnr.2002.8021

66 Lang, A. E. et al. Randomized controlled trial of intraputamenal glial cell line-derived neurotrophic factor infusion in Parkinson disease. Ann Neurol 59, 459–466 (2006). 10.1002/ana.20737

67 Nutt, J. G. et al. Randomized, double-blind trial of glial cell line-derived neurotrophic factor (GDNF) in PD. Neurology 60, 69–73 (2003). 10.1212/wnl.60.1.69

68 Tatarewicz, S. M. et al. Development of a maturing T-cell-mediated immune response in patients with idiopathic Parkinson’s disease receiving r-metHuGDNF via continuous intraputaminal infusion. J Clin Immunol 27, 620–627 (2007). 10.1007/s10875-007-9117-8

69 Slevin, J. T. et al. Improvement of bilateral motor functions in patients with Parkinson disease through the unilateral intraputaminal infusion of glial cell line-derived neurotrophic factor. J Neurosurg 102, 216–222 (2005). 10.3171/jns.2005.102.2.0216

70 Slevin, J. T. et al. Unilateral intraputamenal glial cell line-derived neurotrophic factor in patients with Parkinson disease: response to 1 year of treatment and 1 year of withdrawal. J Neurosurg 106, 614–620 (2007). 10.3171/jns.2007.106.4.614

71 Herzog, C. D. et al. Enhanced neurotrophic distribution, cell signaling and neuroprotection following substantia nigral versus striatal delivery of AAV2-NRTN (CERE-120). Neurobiol Dis 58, 38–48 (2013). 10.1016/j.nbd.2013.04.011

72 Mandel, R. J., Spratt, S. K., Snyder, R. O. & Leff, S. E. Midbrain injection of recombinant adeno-associated virus encoding rat glial cell line-derived neurotrophic factor protects nigral neurons in a progressive 6-hydroxydopamine-induced degeneration model of Parkinson’s disease in rats. Proc Natl Acad Sci U S A 94, 14083–14088 (1997). 10.1073/pnas.94.25.14083

73 Kirik, D., Rosenblad, C., Bjorklund, A. & Mandel, R. J. Long-term rAAV-mediated gene transfer of GDNF in the rat Parkinson’s model: intrastriatal but not intranigral transduction promotes functional regeneration in the lesioned nigrostriatal system. J Neurosci 20, 4686–4700 (2000). 10.1523/JNEUROSCI.20-12-04686.2000

74 Marks, W. J., Jr., et al. Gene delivery of AAV2-neurturin for Parkinson’s disease: a double-blind, randomised, controlled trial. Lancet Neurol 9, 1164–1172 (2010). 10.1016/S1474-4422(10)70254-4

75 Kordower, J. H. et al. Delivery of neurturin by AAV2 (CERE-120)-mediated gene transfer provides structural and functional neuroprotection and neurorestoration in MPTP-treated monkeys. Ann Neurol 60, 706–715 (2006). 10.1002/ana.21032

76 Ruitenberg, M. J., Eggers, R., Boer, G. J. & Verhaagen, J. Adeno-associated viral vectors as agents for gene delivery: application in disorders and trauma of the central nervous system. Methods 28, 182–194 (2002). 10.1016/s1046-2023(02)00222-0

77 Towne, C., Pertin, M., Beggah, A. T., Aebischer, P. & Decosterd, I. Recombinant adeno-associated virus serotype 6 (rAAV2/6)-mediated gene transfer to nociceptive neurons through different routes of delivery. Mol Pain 5, 52 (2009). 10.1186/1744-8069-5-52

78 Bresjanac, M. & Antauer, G. Reactive astrocytes of the quinolinic acid-lesioned rat striatum express GFRalpha1 as well as GDNF in vivo. Exp Neurol 164, 53–59 (2000). 10.1006/exnr.2000.7416

79 Nakagawa, T. & Schwartz, J. P. Gene expression profiles of reactive astrocytes in dopamine-depleted striatum. Brain Pathol 14, 275–280 (2004). 10.1111/j.1750-3639.2004.tb00064.x

80 Yamagata, K. et al. Adenosine induces expression of glial cell line-derived neurotrophic factor (GDNF) in primary rat astrocytes. Neurosci Res 59, 467–474 (2007). 10.1016/j.neures.2007.08.016

81 Azevedo, F. A. et al. Equal numbers of neuronal and nonneuronal cells make the human brain an isometrically scaled-up primate brain. J Comp Neurol 513, 532–541 (2009). 10.1002/cne.21974

82 von Boyen, G. B. et al. Distribution of enteric glia and GDNF during gut inflammation. BMC Gastroenterol 11, 3 (2011). 10.1186/1471-230X-11-3

83 Steinkamp, M. et al. GDNF protects enteric glia from apoptosis: evidence for an autocrine loop. BMC Gastroenterol 12, 6 (2012). 10.1186/1471-230X-12-6

84 Takano, T., Oberheim, N., Cotrina, M. L. & Nedergaard, M. Astrocytes and ischemic injury. Stroke 40, S8–12 (2009). 10.1161/STROKEAHA.108.533166

85 Ouchi, Y. et al. Microglial activation and dopamine terminal loss in early Parkinson’s disease. Ann Neurol 57, 168–175 (2005). 10.1002/ana.20338

86 Rickert, U. et al. Glial Cell Line-Derived Neurotrophic Factor Family Members Reduce Microglial Activation via Inhibiting p38MAPKs-Mediated Inflammatory Responses. J Neurodegener Dis 2014, 369468 (2014). 10.1155/2014/369468

87 Wu, H. M. et al. Novel neuroprotective mechanisms of memantine: increase in neurotrophic factor release from astroglia and anti-inflammation by preventing microglial activation. Neuropsychopharmacology 34, 2344–2357 (2009). 10.1038/npp.2009.64

88 Gomes, C. A., Vaz, S. H., Ribeiro, J. A. & Sebastiao, A. M. Glial cell line-derived neurotrophic factor (GDNF) enhances dopamine release from striatal nerve endings in an adenosine A2A receptor-dependent manner. Brain Res 1113, 129–136 (2006). 10.1016/j.brainres.2006.07.025

89 Kopra, J. J. et al. Dampened Amphetamine-Stimulated Behavior and Altered Dopamine Transporter Function in the Absence of Brain GDNF. J Neurosci 37, 1581–1590 (2017). 10.1523/JNEUROSCI.1673-16.2016

90 Lattanzi, R. et al. Pharmacological activity of a Bv8 analogue modified in position 24. Br J Pharmacol 166, 950–963 (2012). 10.1111/j.1476-5381.2011.01797.x

91 Beale, K. et al. Peripheral administration of prokineticin 2 potently reduces food intake and body weight in mice via the brainstem. Br J Pharmacol 168, 403–410 (2013). 10.1111/j.1476-5381.2012.02191.x

