## Supplementary Information for "Prokineticin-2 Upregulates GDNF in Astrocytes and Pharmacological Modulation of PK2 Receptors offers Neuroprotection in Experimental Models of Parkinson’s Disease"

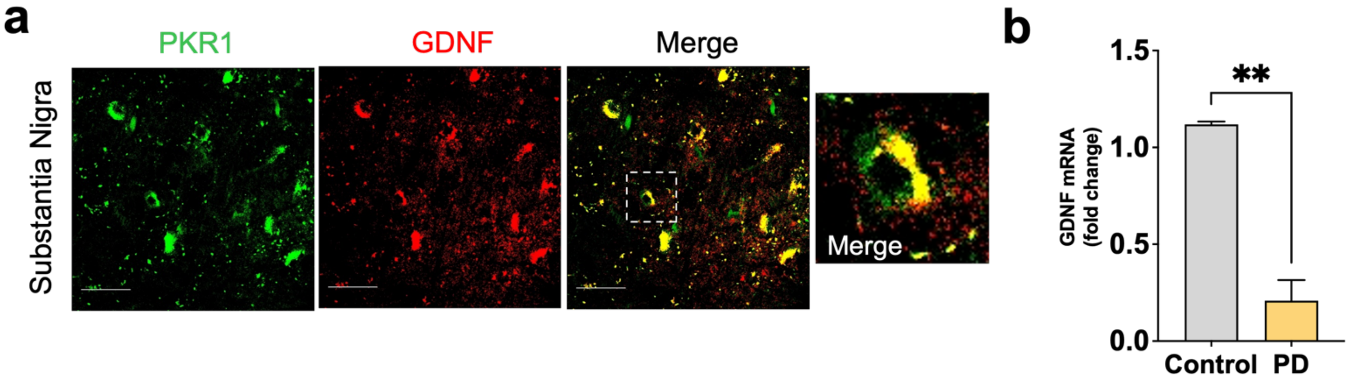


**Supplementary Fig. 1 GDNF expression is reduced in the SN of PD patient samples and colocalizes with PKR1.** (a) Double-labelling immunofluorescence for GDNF (red) and PKR1 (green) in human post-mortem nigral sections. Scale bar = 50 µm. (b) Quantitative PCR showing GDNF downregulation in striatum of PD patients. Data represented as mean ± SEM and expressed as fold-change normalized to control (n=10). **p<0.01 Student's t-test.

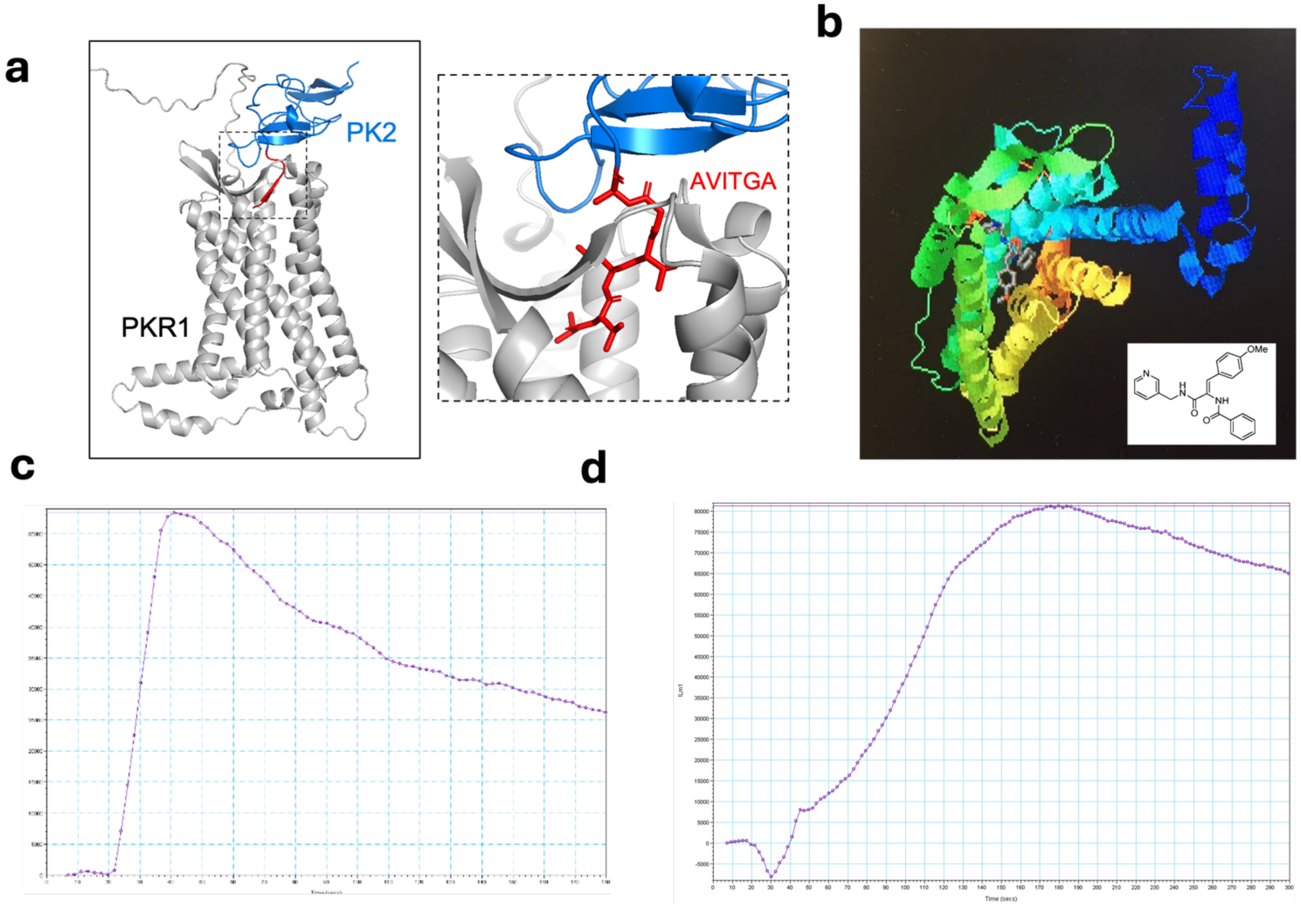


**Supplementary Fig. 2 PK2 and IS20 activate PKR1 and initiate calcium mobilization.** (a) AlphaFold3 modeling of PK2–PKR1 binding with PK2 in blue, the required AVITGA PK2 sequence motif in red, and PKR1 in grey. (b) IS20 complexing with PKR1. (c) Calcium assay analysis of CHO cells stably expressing PKR1 treated with 100 nM rPK2. (d) Calcium assay analysis of CHO cells stably expressing PKR1 treated with 10 μM IS20.

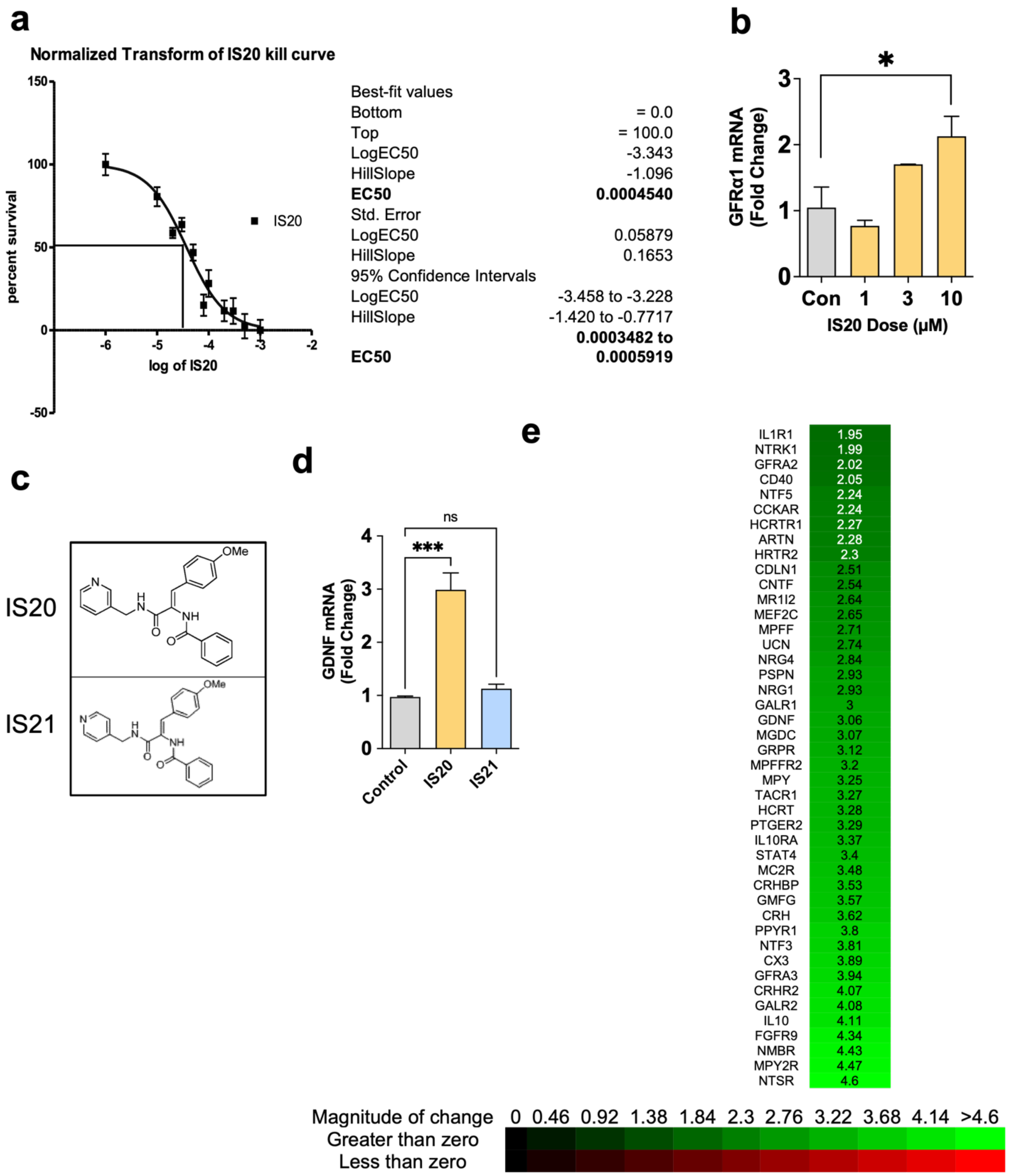


**Supplementary Fig. 3.** (a) Kill curve of primary mouse astrocytes treated with IS20 with corresponding EC50 measurements (right). (b) qPCR analysis of GFRa1 in primary mouse astrocytes treated with (1–10 µM IS20). (c) IS20 and IS21 molecular structures. (d) qPCR analysis of primary mouse astrocytes treated with IS20 (10 µM) or IS21 (10 µM) for 3 h. (e) Neurotrophin gene array analysis of primary mouse astrocytes treated with IS20 (10 µM). *$p \le 0.05$, $p < 0.001$.
